# Template strand deoxyuridine promoter recognition by a viral RNA polymerase

**DOI:** 10.1101/2021.08.30.458274

**Authors:** Alec Fraser, Maria L. Sokolova, Arina V. Drobysheva, Julia V. Gordeeva, Sergei Borukhov, Tatyana O. Artamonova, AlphaFold team, Konstantin V. Severinov, Petr G. Leiman

## Abstract

*Bacillus subtilis* bacteriophage AR9 employs two strategies for efficient host takeover control and host antiviral defense evasion – it encodes two unique DNA-dependent RNA polymerases (RNAPs) that function at different stages of virus morphogenesis in the cell, and its double stranded (ds) DNA genome contains uracils instead of thymines throughout^1,2^. Unlike any known RNAP, the AR9 non-virion RNAP (nvRNAP), which transcribes late phage genes, recognizes promoters in the template strand of dsDNA and efficiently differentiates obligatory uracils from thymines in its promoters^3^. Here, using structural data obtained by cryo-electron microscopy and X-ray crystallography on the AR9 nvRNAP core, holoenzyme, and a promoter complex, and a variety of *in vitro* transcription assays, we elucidate a unique mode of uracil-specific, template strand-dependent promoter recognition. It is achieved by a tripartite interaction between the promoter specificity subunit, the core enzyme, and DNA adopting a unique conformation. We also show that interaction with the non-template strand plays a critical role in the process of AR9 nvRNAP promoter recognition in dsDNA, and that the AR9 nvRNAP core and a part of its promoter specificity subunit that interacts with the core are structurally similar to their bacterial RNAP counterparts. Our work demonstrates the extent to which viruses can evolve new functional mechanisms to control acquired multisubunit cellular enzymes and make these enzymes serve their needs.

---

*Bacillus subtilis* “jumbo” bacteriophage AR9 encodes two distinct multisubunit DNA-dependent RNA polymerases (RNAPs), allowing for the transcription of viral genes to proceed independently of the host RNAP^1–4^. The virion-packaged RNAP (vRNAP) is delivered into the host cell together with phage DNA at the onset of infection. The vRNAP then transcribes early phage genes, including those of the second, non-virion RNAP (nvRNAP). The nvRNAP transcribes late genes, including those coding for the vRNAP, which is then packaged into progeny phage particles.

The catalytically active AR9 nvRNAP core enzyme consists of four proteins that, when pairwise concatenated, show about 20% sequence identity and cover the entire length of the universally conserved β and β′ subunits of bacterial RNAPs (**Extended data Fig. 1a**)^3^. Promoter-specific transcription is performed by a five-subunit holoenzyme that in addition to the nvRNAP core contains the product of AR9 gene *226* (gp226)^3^. Gp226 shows no sequence similarity to bacterial RNAP σ subunits which mediate the recognition of promoters in bacteria or any known transcription factor. Close orthologs of gp226 are found in the genomes of other jumbo phages that have been demonstrated or are presumed to contain uracil in their genomic DNA^5,6^.

Unlike bacterial RNAPs^7^, the AR9 nvRNAP holoenzyme recognizes promoters in the template strand of dsDNA and is capable of promoter-specific transcription initiation on single stranded (ss) DNA^3^. The AR9 nvRNAP promoter consensus 3′-^−11^UUGU^−8^-N_6_-AU^+1^-5′ (where N is any nucleotide and the transcription start site (TSS) coordinate is +1) contains a four-base long motif centered about 10 bases upstream of the TSS and two bases at the TSS (**Extended data Fig. 1b**). Promoters with thymines in the −11^th^ and −10^th^ positions are inactive suggesting that the C5 position of the uracil’s pyrimidine ring, which carries a methyl group in the thymine, plays a critical role in promoter recognition (**Extended data Fig. 1c**). Despite possessing a short promoter consensus element, the AR9 nvRNAP holoenzyme protects an extensive region of DNA flanking the TSS (position −35 to +20 in the template strand and positions −29 to +17 in the non-template strand) from DNase I attack, implying additional contacts with DNA^3^.

To understand the uracil-specific, template strand-dependent promoter recognition mechanism of the AR9 nvRNAP, we determined the structure of this enzyme by X-ray crystallography and cryo-electron microscopy (cryo-EM) in three states – the core, holoenzyme, and holoenzyme in complex with a 3′-overhang dsDNA oligonucleotide (historically called a “forked” or “fork” template) that mimicked one half of the transcription bubble (**Fig. 1a**). Furthermore, we complemented this structural information by discriminative *in vitro* transcription assays. The 18 base-long ss part of the dsDNA oligonucleotide contained the late AR9 promoter P077 while its 14 bp-long ds segment spanned positions from +3 to +16 relative to the TSS (**Fig. 1b**). Fortuitously, in the most populous class of the cryo-EM reconstruction and in both available crystal forms, the enzyme bound not one but two copies of this oligonucleotide – a downstream copy, as designed, and an upstream one, unexpectedly – resulting in a superstructure that resembled the complete transcription bubble found in open complexes formed by other RNAPs. Moreover, *in crystallo* the nvRNAP molecules and the oligonucleotides formed a train in which the upstream and downstream oligonucleotides belonging to two neighboring unit cells pi-stacked and formed a continuous double helix (**Extended Data Fig. 2**).

**Fig. 1.**
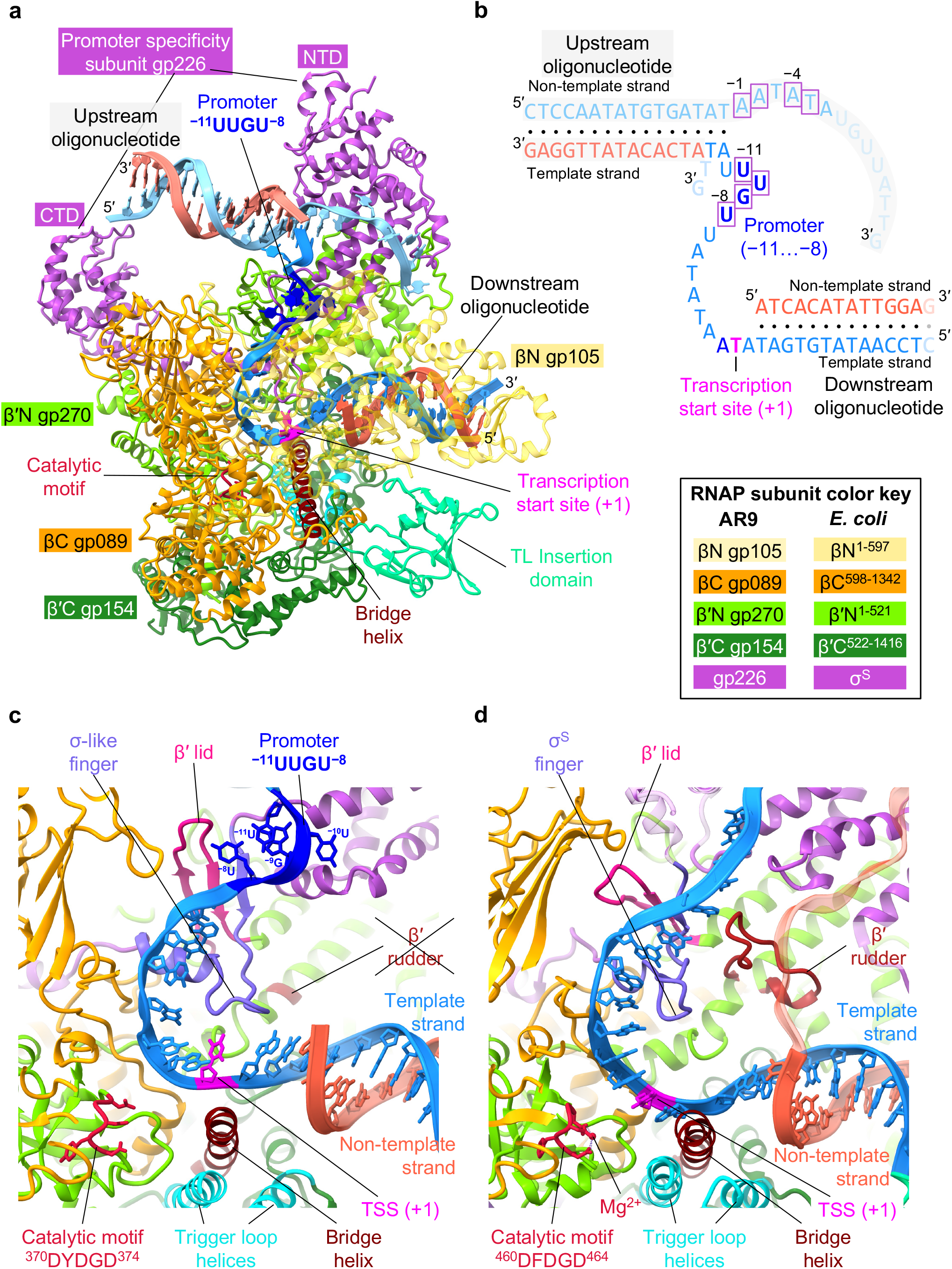
Structure of the AR9 nvRNAP promoter complex. **a**, Ribbon diagram of the crystal structure of the AR9 nvRNAP in complex with a 3′-overhang dsDNA (fork) oligonucleotide. Structural elements that are either unique to the AR9 nvRNAP or common to all RNAPs are labeled and color coded. The βN gp105 subunit is semitransparent for clarity. **b**, Schematic of the two oligonucleotides that bound to one AR9 nvRNAP molecule resulting in a transcription bubble-like structure. Bases disordered in the crystal structure are rendered semitransparent. Bases in purple boxes interact with the protein. **c** and **d**, Structure of the catalytic centers of the AR9 nvRNAP and *E. coli* RNAP-σ^S^ (PDB code 5IPM^15^). Here and elsewhere, TSS stands for the transcription start site. The 2.4 region of σ^S^, which is not present in gp226, is rendered semi-transparent. The *E. coli* RNAP-σ^S^ structure contains a short RNA product that is not shown for clarity. A part of the DNA non-template strand in the *E. coli* RNAP-σ^S^ structure is semitransparent for clarity.

## Structural similarities and differences between the AR9 nvRNAP and bacterial RNAPs

The overall structure of the AR9 nvRNAP is a trimmed down version of a bacterial crab claw-shaped RNAP (**Fig. 1a**, **Extended Data Fig. 3**). No domain compensates for the absence of α and ω subunits that are present in all bacterial, eukaryotic, and archaeal enzymes. As a result, the AR9 nvRNAP claw is smaller and has a boxier appearance than that of its cellular counterparts. In bacterial enzymes, the α subunit dimer serves as a platform for the assembly of the β and β′ subunits^8–10^. In the AR9 nvRNAP structure, the split site of the β′ subunit is spatially close to the putative β′-α^II^ interface (**Extended data Fig. 3**), which suggests that this location likely represents a critical point for the formation of tertiary and quaternary structure. By splitting its β and β′ subunits into smaller proteins, the AR9 nvRNAP appears to have removed this assembly requirement.

Inside the catalytic cleft, the AR9 nvRNAP core contains all of the structural elements required for catalysis, stabilization of the open promoter complex, and promoter clearance found in multisubunit RNAPs^9,11^, except for the β′ rudder (**Fig. 1c, 1d, Extended data Fig. 3**). The β′ rudder is a twisted β-hairpin that extends from one of the β′ clamp α-helices and interacts with the RNA-DNA hybrid near the active site^9^. In bacterial RNAPs, its deletion impairs promoter opening and destabilizes the elongation complex but does not affect the efficiency of transcription termination or the length of the RNA-DNA hybrid^12^. The elongation complex of the AR9 nvRNAP is likely stabilized by a different mechanism. The conformation of the ^370^DYDGD^374^ catalytic motif of the AR9 nvRNAP, which is located near the C terminus of the β′N subunit gp270, is similar to that found in other RNAPs. The side chains of the three conserved aspartates are poised to bind a Mg^2+^ ion that is universally conserved in all nucleotidyltransferases, albeit the resolution of X-ray and cryo-EM data is insufficient for resolving it (**Fig. 1c, 1d, Extended data Fig. 3a**).

Similar to the *E. coli* RNAP^13^, the AR9 nvRNAP contains an insertion domain (residues 400-508 of β′C gp154) in the trigger loop (**Fig. 1a, Extended data Fig. 3**). The trigger loop undergoes major conformational changes during the catalytic nucleotide addition cycle and template translocation^9,11^, and the presence of an insertion domain in the *E. coli* system has not been fully reconciled with these transformations^14^. The fold of the AR9 nvRNAP insertion domain is different from that of the *E. coli* RNAP and, in fact, to any protein in the Protein Data Bank (PDB). Its sequence is also unique and found only in nvRNAPs of other jumbo phages^5,6^. Its position in the structure of the nvRNAP core is also different from that of the *E. coli* RNAP.

In the AR9 nvRNAP core structure, the insertion domain is located roughly in-between the β and β′ pincers where it partially obstructs the downstream DNA channel (**Extended data Fig. 4a**). Furthermore, it carries a strong negative charge on its DNA-facing surface suggesting that it interacts with the downstream dsDNA in a non-sequence specific manner (**Extended data Fig. 4b**). Out of 10 independent copies of the AR9 nvRNAP core molecules belonging to two different crystal forms, the insertion domain is ordered in only a singular instance. In the cryo-EM structure of DNA template-free holoenzyme, this domain is fully disordered (**Extended data Fig. 5a, 5b**). Considering the intrinsic propensity of this domain to large motions, it may participate in translocation by sliding on the DNA and exerting a force on the trigger loop.

## The structure of promoter-specificity subunit gp226

The AR9 nvRNAP promoter-specificity subunit gp226 consists of two globular domains – a larger N-terminal domain (NTD, residues 1-264) and a smaller C-terminal domain (CTD, residues 295-464) – connected by a linker (**Fig. 1a, Extended data Fig. 6a**).

Gp226 interacts with the nvRNAP core in a manner resembling that of bacterial σ factors ^7,11,15,16^, and all elements that come in contact with the body of the enzyme have structural counterparts in bacterial σ factors. Residues 184-264 of gp226 fold into a σ_2_-like domain (**Extended data Fig. 6a**), which interacts in a sequence specific manner with the non-template strand of the −10 promoter element in bacterial σ factors and comprises their most conserved part^17–19^ (**Extended data Fig. 6b**). Gp226 residues 265-294 form a σ finger-like structure that invades the catalytic cleft and forms an augmented β-sheet with the β′ lid (**Fig. 1c**, **Extended data Fig. 6a**). Gp226 residues 295-316 comprise an α-helix that matches the N-terminal α-helix of the σ_4_ domain (**Extended data Fig. 6a, 6b**).

The overall structural and sequence similarities of gp226 to bacterial σ factors are low. The most similar σ factor with a known structure, the *E. coli* σ^E^, displays a Cα-Cα root mean square deviation (RMSD) of 4.2 Å and a sequence identity of 9.6% when superimposed onto residues 184-316 of gp226 which comprise its σ_2_-, finger- and σ_4_-like elements. Nevertheless, considering that i) the σ-like part of gp226 is nearly equal in size to the entire σ^E^ structure (**Extended data Fig. 6a, 6b**), ii) this part interacts with the core enzyme over an extended distance, and iii) the gp226 transcription bubble structure resembles that of bacterial RNAP-σ holoenzymes, it follows that gp226 is likely a homolog of bacterial σ factors in which the peripheral parts of the N- and C-terminal domains have been replaced with new folds but all elements that interact with the RNAP core have been retained.

A Protein Data Bank-wide search^20^ for folds resembling that of the gp226 NTD and CTD resulted in a single definitive match. The CTD of gp68, a subunit of the phage phiKZ non-virion RNAP which is required for elongation and promoter recognition^21,22^, can be superimposed onto the gp226 CTD with an RMSD of 3.2 Å for 142 equivalent Cα atoms (out of 173) and a sequence identity of 11% (**Supplementary Fig. 2a**). As the rest of the gp68 structure (residues 1-303) is disordered, we used AlphaFold^23^ Colab to model it. The local distance difference test^24^ of this model for residues 1-277 was 85.3, indicating a very high level of confidence (**Supplementary Fig. 2b**). This model can be superimposed onto the gp226 NTD with an RMSD of 3.0 Å for 198 equivalent Cα atoms (out of 277) and a sequence identity of 8.6% (**Supplementary Fig. 2a**). Thus, even though gp226 is divergent from gp68 sequence-wise, their similarities in structure and locations within the RNAP holoenzyme complex suggest that the two proteins have a common ancestor.

## Structural adaptations of gp226 required for template strand promoter recognition

The most conserved parts of bacterial σ factors, the σ_2_- and σ finger elements, are also conserved in gp226 (**Extended data Fig. 6**). These elements, with small adaptations, are responsible for the recognition of the promoter in the template strand of DNA. A tight turn connecting helices 2.1 and 2.2 in bacterial σ factors is replaced by a short α-helix in gp226 (residues 205-214). This 2.1bis helix creates a bridge linking the two pincers of the AR9 nvRNAP claw and together with the βN gp105 subunit comprises a binding site for the promoter in the template strand of DNA (**Extended data Fig. 6a**). The gp226 finger forms an augmented β sheet with the β′ lid (residues 161-177 of β′N gp270) such that the β′ lid reaches the −8 position uracil base of the 3′-^−11^UUGU^−8^-5′ promoter motif and tucks it in against the 2.1bix helix (**Fig. 1c, 1d**). The β′ lid of AR9 nvRNAP is three residues longer than its bacterial counterpart, which further enhances and facilitates this interaction (**Fig. 1c**).

## Promoter DNA structure and the design of a T-specific enzyme

The structure of the 3′-^−11^UUGU^−8^-5′ template strand promoter motif is well resolved in both cryo-EM and X-ray electron density maps, although ^−8^U is partially disordered in the cryo-EM map (**Extended data Fig. 7a, 7b).** The most critical and obligatory ^−10^U base, the replacement of which by a T leads to the abolishment of promoter-specific transcription^3^ (**Extended Data Fig. 1c**), is buried in a deep pocket at the interface of the gp226 2.1bis helix and βN gp105 (**Fig. 2a**). In this pocket, the ^−10^U base is wedged between the side chains of gp226 I207 and βN gp105 R363, forming a stacking interaction with the latter. Its Watson-Crick interface forms hydrogen bonds with the side chain of βN gp105 K375 and with the main chain N of gp226 V206 and I207. Most importantly, the C5 atom of the ^−10^U pyrimidine ring is only 3.9 Å away from the Cβ of V206, suggesting that a C5 position methyl group would clash with the V206 side chain (**Fig. 2a**). Accordingly, a V206G mutant recognized ^−10^U- and ^−10^T-containing promoters with equal efficiencies (**Fig. 2b**). Notably, all close homologs of gp226 proteins in jumbo phages with deoxyuridine-containing genomic DNA^5,6^ display high sequence conservation of the 2.1bis helix with the critical valine being absolutely conserved. This suggests that these phages employ a similar mechanism for uracil-dependent promoter recognition.

**Fig. 2.**
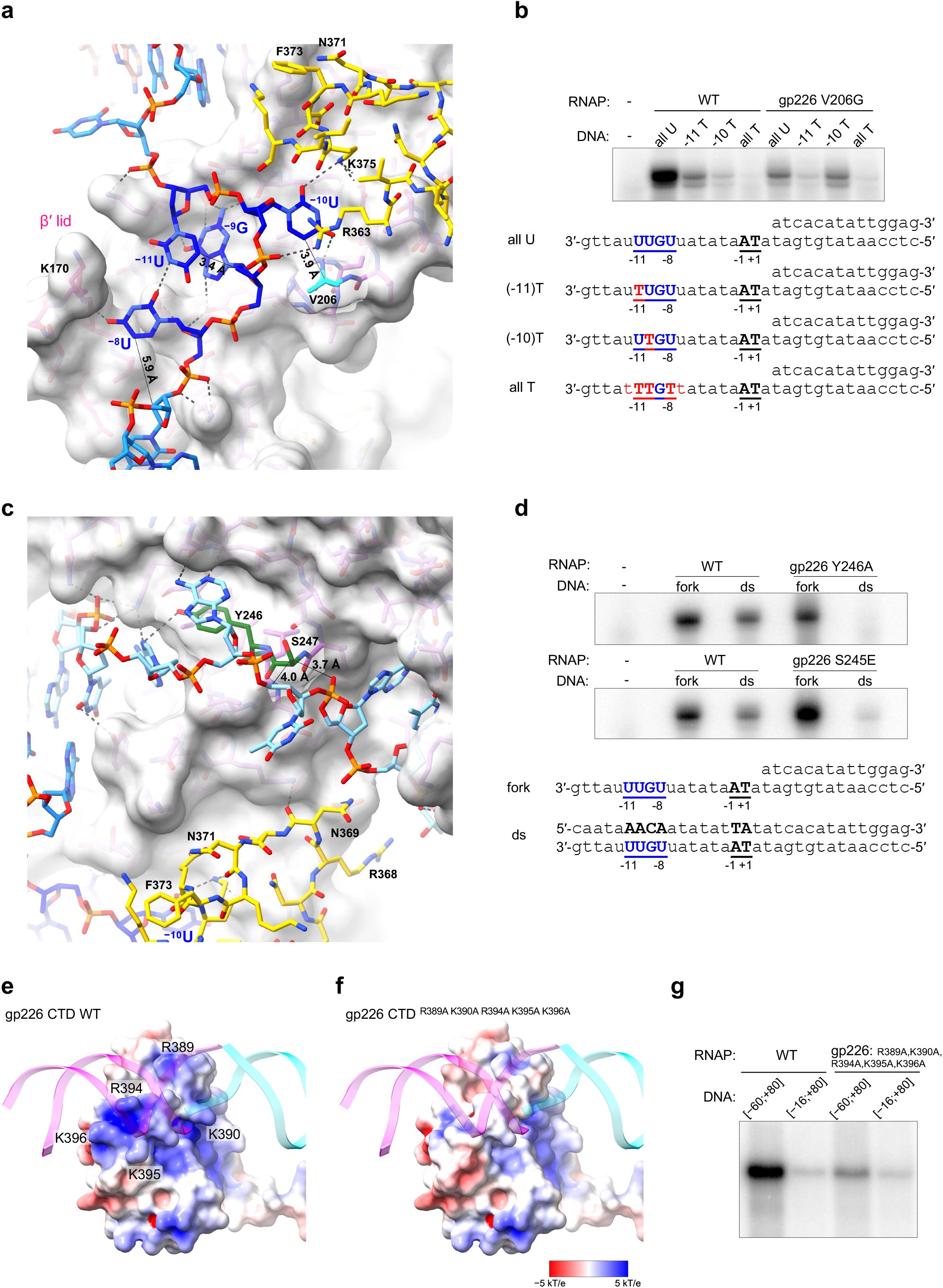
Interaction of the AR9 nvRNAP with DNA in the promoter complex. **a**, Atomic model of the AR9 nvRNAP promoter recognition element. Gp226 is shown as a semitransparent molecular surface. Only a small fragment of βN gp105 that participates in the formation of the ^−10^U binding pocket is shown for clarity. Interchain and DNA-intrachain hydrogen bonds are shown as dashed lines. Thin, straight lines connect the C5 atom of the uracil pyrimidine ring to the closest protein or DNA atoms which lie in-plane with the ring. The carbon atoms are colored as in **Fig. 1a**, except for V206, which is shown in cyan. **b**, The *in vitro* transcription activity of the AR9 nvRNAP holoenzyme containing the wild type (WT) gp226 or V206G gp226 mutant have been tested using various U- and T-containing templates. **c**, The non-template strand forms extensive interactions with the gp226 NTD. The same gp226 and βN gp105 fragments as in **Fig. 2a** are shown but are tilted to improve clarity. **d**, The *in vitro* transcription activity of AR9 nvRNAP mutants with an altered structure of the non-template strand binding groove. **e** and **f**, The surface electrostatic potentials of the pseudo^−35^-element binding motif in the WT gp226 and A^5^ gp226 mutant. **g**, The ssDNA and dsDNA *in vitro* transcription activities of the AR9 nvRNAP WT and A^5^ mutant. For each *in vitro* transcription experiment, two technical replicates of two biological replicates resulted in similar outcomes and one of them is shown. The uncropped autoradiograms are presented in **Supplementary Fig. 1**.

The requirement of U vs. T in the −11^th^ position of the promoter is nearly as strong as in the −10^th^ position. Additionally, G is required in the −9^th^ position^3^. However, the enzyme displays almost no U vs. T preference in the −8^th^ position (**Extended data Fig. 1c)**. In the promoter complex, the bases of ^−11^U and ^−9^G form a stacking interaction such that the C5 and C6 atoms of ^−11^U butt against the sugar-phosphate backbone of the ^−11^UUG^−9^ segment, leaving no space for a C5 position methyl group (**Fig. 2a**). There is a stacking interaction between ^−9^G and the phenol ring of gp226 Y210 (which belongs to the 2.1bis helix), and there are three hydrogen bonds between the Watson-Crick interface of ^−9^G and the main chain of gp226 residues F261 and Y263, which provides a rationale for the G requirement in this position. ^−8^U forms one hydrogen bond with ^−11^U and one hydrogen bond with the tip of the β′ lid, which is longer than in its bacterial counterparts, as described above. The C5 position of ^−8^U points into solution and can accommodate the additional methyl group of T (**Fig. 2a**).

## Interaction with the non-template strand is essential for the transcription of dsDNA

The gp226 NTD displays several deep pockets that capture the ss part of the upstream oligonucleotide, which mimics the non-template strand of the transcription bubble (**Fig. 2c**). Considering the structure of the transcription bubble, this part of the non-template strand must be complementary to the template strand promoter, that is its sequence should be 5′-^−11^AACA^−8^-3′ where +1 is the TSS. Our oligonucleotide contained a similar motif in its ss part (5′-^−1^AATA^−4^-3′, **Fig. 1b**). Together with its neighboring bases this sequence matched the appearance of the electron density. This motif was likely a key determinant in the fortuitous binding of the upstream oligonucleotide.

The gp226 NTD interacts with the backbone and bases of the non-template DNA strand of the transcription bubble via pi-pi stacking, ion pairs, and hydrogen bonds. The length of this interface exceeds 30 Å. The extent of these interactions suggests that they play an important role in promoter recognition of a dsDNA template. Indeed, a Y246A mutation that eliminates pi-pi stacking between the side chain of Y246 and the ^−2^A base of the ^−1^AATA^−4^ motif abolishes transcription on dsDNA but does not affect transcription on a fork template (**Fig. 2d**). Similarly, a S245E mutation introduces a large, negatively charged side chain that interferes with the trace and conformation of the sugar-phosphate backbone between ^−2^A and ^−4^A on the surface of the gp226 NTD. As a consequence, the transcriptional activity of the S245E mutants on a dsDNA template is weak, whereas its fork template activity is at or above that of WT (**Fig. 2d**).

In the cryo-EM structure of the DNA template-free AR9 nvRNAP holoenzyme, the NTD and σ-like finger of gp226 are disordered (**Extended data Fig. 5a, 5b**). Similarly to the TL insertion domain, the disorder is likely due to positional heterogeneity since both the gp226 NTD and TL insertion domain possess well-defined hydrophobic cores and are folded in other states of the nvRNAP complex. Furthermore, the NTD is resistant to proteolysis by trypsin in gp226 recombinantly expressed on its own (**Extended Data Fig. 1d**). The order-disorder transitions of the gp226 NTD and σ-like finger play a role in the promoter recognition mechanism described below.

## Gp226 CTD interacts with DNA in a non-sequence specific manner

Although weak, the cryo-EM and X-ray electron density of the upstream oligonucleotide stretches from the σ2-like part of the gp226 NTD to the gp226 CTD. This interaction is about 35 DNA bases upstream from the TSS (**Extended data Fig. 6a**) drawing a parallel to the −35 consensus element of bipartite bacterial promoters. The recognition of the −35 element by bacterial σ factors is mediated by a helix-turn-helix motif^25^, which interacts with the major groove of dsDNA^26^ in a sequence specific manner (**Extended data Fig. 6b**). AR9 nvRNAP promoters, however, display no sequence conservation in this region (**Extended data Fig. 1b**) and, accordingly, the gp226 CTD interacts with the minor groove of dsDNA which displays few sequence-specific features in the B form^27^. Furthermore, this interaction is mediated by a scrunched β-strand that carries several positively charged residues (R389, K390, R394, K395, K396), but not by a helix-turn-helix motif, which is absent from the gp226 structure. We termed the DNA interacting element of the gp226 CTD (amino acids 386-395) a pseudo-^−35^element-binding motif.

To examine the role of the pseudo-^−35^element-binding motif in promoter recognition, we removed most of the positive charge displayed on its surface by mutating R389, K390, R394, K395, K396 of gp226 to alanines (we called this mutant A^5^) (**Fig. 2e, 2f**). As the A^5^ mutant had a lower activity overall, we compared its activity to that of the wild type (WT) on two dsDNA templates that either contained or lacked an upstream part required for interaction with the pseudo-^−35^element-binding motif (the [−60,+80] and [−16,+80] templates in **Fig. 2g**, respectively). The WT enzyme exhibited a greater decrease in activity on the [−16,+80] template compared to that of the A^5^ mutant, which shows that the pseudo-^−35^element-binding motif is essential for optimal promoter recognition.

The AR9 gp226 pseudo-^−35^element-binding motif maps onto a disordered part of phage phiKZ gp68^21^ (residues 413-429, **Supplementary Fig. 2a**). This region of gp68’s surface carries a positive charge, akin to that of AR9 gp226 (**Fig. 2e**), even without the inclusion of disordered residues in electrostatic potential calculations. Again, analogous to the AR9 nvRNAP, phiKZ nvRNAP promoters show no sequence conservation 35 bases upstream of the TSS^28^. Considering i) the homology of the phiKZ nvRNAP to cellular RNAPs^21^ and, by extension, to the AR9 nvRNAP, ii) the homology of AR9 gp226 to phiKZ gp68 described above, and iii) the presence of positive charges at equivalent locations on the surface of the AR9 gp226 and phiKZ gp68 CTDs, the phiKZ nvRNAP is thus likely to form an AR9 nvRNAP-like transcription bubble in which the CTD of gp68 participates in the binding of upstream dsDNA. Furthermore, as all these properties are seemingly conserved for such distantly related viruses as AR9 and phiKZ that i) infect unrelated hosts (Gram positive *B. subtilis* and Gram negative *Pseudomonas aeruginosa*, respectively), ii) encode proteins that show nearly random levels of sequence identity, and iii) have different genomic DNA base compositions^1,29^, the supposition of AR9 nvRNAP-like transcription bubbles can be extended to nvRNAPs of all jumbo phages.

## Free energy of template strand promoter binding

To reconcile the tight integration of promoter DNA into the promoter complex with the transient nature of this complex (**Fig. 1a, 1c, 2a**) we examined the binding free energy of the three best-ordered promoter bases 3′-^−11^UUG^−9^-5′ to the AR9 nvRNAP holoenzyme by executing a double decoupling method molecular dynamics protocol ^30–33^. The procedure assumes that the conformation of the enzyme does not appreciably change upon promoter binding. As such, the simulations describe a state in which the gp226 NTD has associated with the AR9 nvRNAP core.

The standard binding free energy was calculated by combining the results of four separate simulations that corresponded to the vertical reactions in the thermodynamic cycle shown in **Extended data Fig. 8a**. After the equilibration of the system (**Extended data Fig. 8b, 8c**), two types of simulations were performed: i) “alchemical transformations” in which the occupancy of the oligonucleotide located either in the promoter pocket 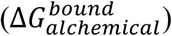 or in bulk water 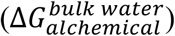 was reduced to zero while the oligonucleotide was harmonically constrained to maintain the promoter pocket bound conformation (**Extended data Fig. 8d, 8e**), and ii) calculations of the entropic cost of such harmonic constraints for a oligonucleotide located in the promoter pocket 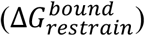 (**Extended data Fig. 8f-8l**) and in bulk water 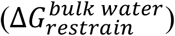 (**Extended data Fig. 8m**). To ensure reproducibility and to minimize bias, all simulations were run bidirectionally.

The favorable energetics of promoter binding via alchemical transformation (−12.7±2.3 kcal/mol, **Extended data Fig. 8d, 8e**) are partially offset by the unfavorable entropic contributions of constraints on DNA conformation and position (5.8±1.5 kcal/mol, **Extended data Fig. 8f-8m**). The resulting free energy gain upon complex formation is −6.9±2.8 kcal/mol, which shows that the interaction of this promoter element with the enzyme is fairly weak. Thus, despite its unusual structure in which the ^−10^U is buried in a deep pocket and ^−9^G and ^−11^U form a stacking interaction, the promoter complex is transient, and the enzyme can easily proceed towards elongation.

## The mechanism of template strand promoter recognition in dsDNA

Combining these findings, we propose the following model for promoter recognition by the AR9 nvRNAP (**Fig. 3**). In the free state of the AR9 nvRNAP holoenzyme molecule, the NTD of gp226 is folded but does not interact with the body of the enzyme (it is positionally disordered or mobile) and the promoter-binding pocket is absent (**Extended view Fig. 5**). The NTD of gp226 is attached to the CTD and the core via a linker that will eventually form a σ finger-like structure in the promoter complex. The enzyme displays two positively charged patches that have DNA binding propensity on their surface – the pseudo-^−35^element-binding motif and a patch with a much stronger positive charge and better shape complementarity for the binding of a non-template strand motif that is complementary to the promoter (**Extended data Fig. 9** and **State 0 in Fig. 3**).

**Fig. 3.**
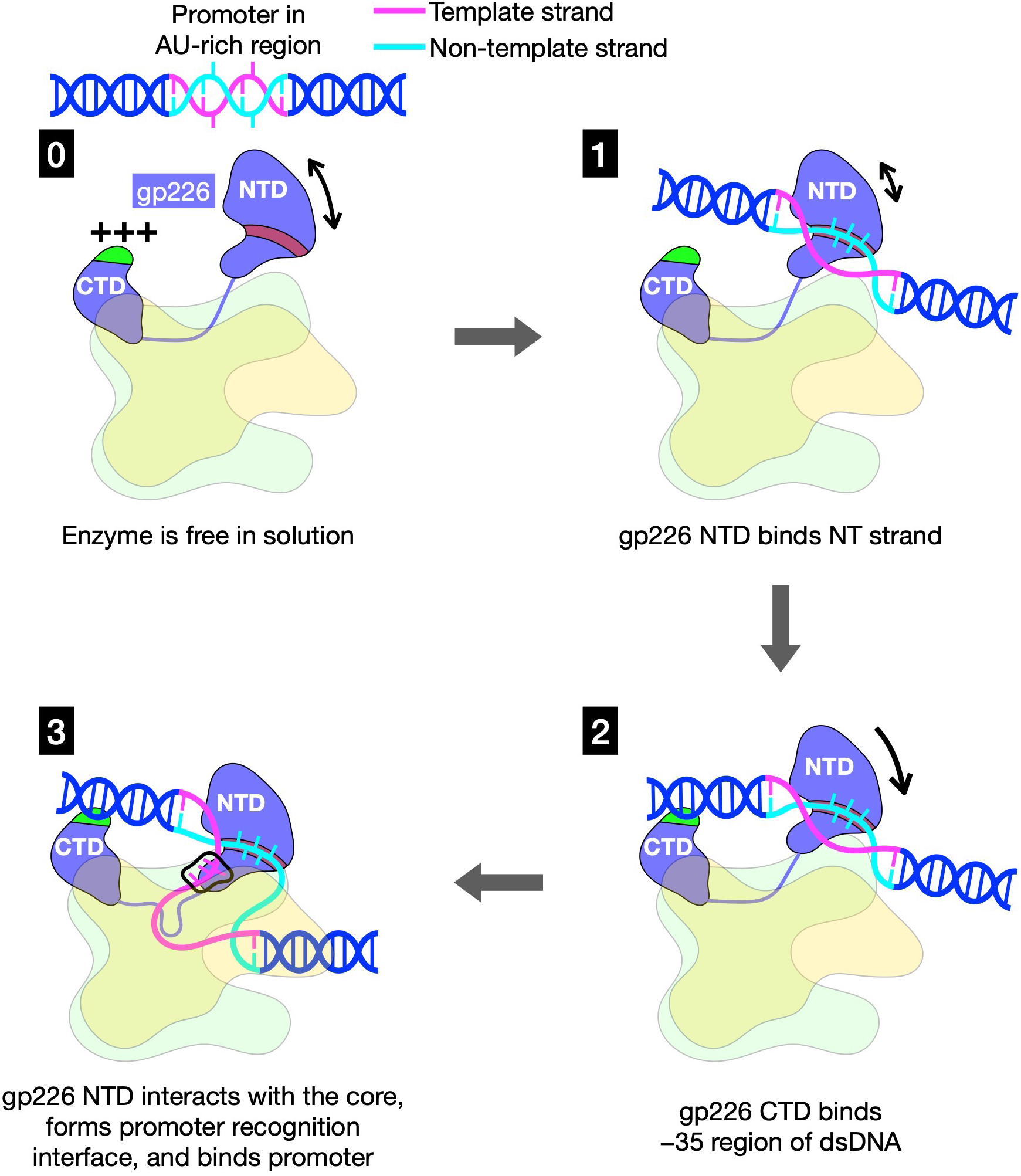
The mechanism of template strand promoter recognition in dsDNA. See the main text for details. For clarity, both proteins comprising the N- and C-terminal parts of the β and β′ subunits are shown in the same color (light yellow for β and light green for β′).

The process of promoter recognition contains the following steps: 1) the gp226 NTD captures an A-rich, promoter-complementary sequence of the non-template strand in a groove on its surface and partially melts dsDNA (**State 1 in Fig. 3**); 2) the pseudo-^−35^element-binding motif of the gp226 CTD interacts with dsDNA, reducing the conformational space available to the gp226 NTD and promoting its binding to the body of the enzyme (**State 2 in Fig. 3**); 3) the NTD of gp226 comes in contact with the body of the enzyme, fully separating the DNA strands, creating a transcription bubble, and placing the template strand at the [gp226]:[βN gp105] interface; this interaction creates a three-dimensional promoter recognition interface that captures a flipped out ^−10^U base and buries it into the ^−10^U recognition pocket while simultaneously squeezing the DNA strand slightly such that the bases that flank the flipped-out ^−10^U base form a stack and the identities of the stacked ^−9^G and ^−11^U bases are verified via geometry-sensitive interactions (hydrogen bonds and ion pairs) (**State 3 in Fig. 3**). As the free energy of promoter recognition is nevertheless reasonably low and the conformation of the sugar-phosphate backbone for the four bases of the promoter motif is close to that of dsDNA, the enzyme can efficiently proceed with elongation.

## Conclusion

Here we have explained the functional mechanism of a phage-encoded RNAP that contains a unique promoter-specificity subunit, recognizes the promoter in the template strand of DNA, requires uracil bases in the promoter, and does not use an otherwise universally conserved helix-turn-helix motif for the binding of dsDNA. Despite its apparent rarity, such enzymes are widespread in the biosphere as the number of such bacteriophages is astronomical. The scarcity of similar sequences is only due to the difficulty of sequencing genomes containing modified nucleotides^5,34^. The extent to which the AR9 nvRNAP promoter recognition mechanism is different from any known RNAP shows that our knowledge about the structure and function of these nanomachines is far from complete.

## Methods

### Cloning of the AR9 nvRNAP and its mutants

Four gene-Blocks (gBlocks) encoding AR9 nvRNAP core enzyme genes optimized for expression in *E. coli* were synthetized by Integrated DNA Technologies (IDT). These gBlocks were assembled into an expression vector on the pETDuet-1 plasmid backbone with the help of the NEBuilder HiFi DNA Assembly Master Mix (New England Biolabs). First, pETDuet-1 was digested by the NcoI and BamHI endonucleases, and gBlocks coding for N-terminally hexahistidine-tagged gp270 and gp154 were ligated. Then, this plasmid was digested by the BglII and XhoI endonucleases and ligated with two gBlocks coding for gp105 and gp089. The resulting plasmid encoded the AR9 nvRNAP core enzyme.

The plasmid for expression of the AR9 nvRNAP holoenzyme was created by inserting an *E. coli*-optimized gp226 gBlock (also synthetized by ITD) into the AR9 nvRNAP core plasmid described above, which was linearized at the XhoI site. This plasmid was used as a template to create mutant versions of the AR9 nvRNAP holoenzyme by site-directed mutagenesis (the list of corresponding primers is in the **Supplementary Table 1**).

The plasmid encoding the tagless AR9 nvRNAP core enzyme was derived from the His-tagged AR9 nvRNAP core plasmid described above. First, a fragment that contained all the four genes but excluded the Hig-tag was PCR amplified (the primers are listed in the **Supplementary Table 1**). Then, the pETDuet-1 vector was linearized by the NcoI and XhoI endonucleases. A new plasmid was then created by ligating the PCR fragment and the linearized pETDuet-1 vector using the NEBuilder HiFi DNA Assembly Master Mix (New England Biolabs).

In all plasmids, a T7 RNAP promoter, a *lac* operator, and a ribosome binding site were located at appropriate positions upstream of each gene.

### Purification of recombinant AR9 nvRNAP

Plasmids encoding AR9 nvRNAP core, tagless AR9 nvRNAP core, and holoenzyme or its mutants were transformed into BL21 Star (DE3) chemically competent *E. coli* cells. The cultures (3 L) were grown at 37°C to OD600 of 0.7 in LB medium supplemented with ampicillin at a concentration of 100 μg/mL, and recombinant protein overexpression was induced with 1 mM IPTG for 4 hours.

Cells containing over-expressed AR9 nvRNAP holoenzyme or its mutants were harvested by centrifugation and disrupted by sonication in buffer A (40 mM Tris-HCl pH 8, 300 mM NaCl, 3 mM β-mercaptoethanol) followed by centrifugation at 15,000 g for 30 min. Cleared lysate was loaded onto a 5 mL HisTrap sepharose HP column (GE Healthcare) equilibrated with buffer A. The column was washed with buffer A supplemented with 20 mM Imidazole. The protein was eluted with a linear 0-0.5 M Imidazole gradient in buffer A. Fractions containing AR9 nvRNAP holoenzyme or its mutants were combined and diluted with buffer B (40 mM Tris-HCl pH 8, 0.5 mM EDTA, 1 mM DTT, 5% glycerol) to the 50 mM NaCl final concentration and loaded on equilibrated 5 mL HiTrap Heparin HP sepharose column (GE Healthcare). The protein was eluted with a linear 0-1 M NaCl gradient in buffer B. Fractions containing AR9 nvRNAP holoenzyme or its mutants were pooled and concentrated (Amicon Ultra-4 Centrifugal Filter Unit with Ultracel-50 membrane, EMD Millipore) to a final concentration of 3 mg/mL, then glycerol was added up to 50% to the sample for storage at −20°C (the samples were used for transcription assays).

Samples used for crystallization and cryo-EM were produced by following a slightly different procedure. Cells containing over-expressed recombinant AR9 nvRNAP core or holoenzyme were harvested by centrifugation and disrupted by sonication in buffer C (50 mM NaH_2_PO_4_ pH 8, 300 mM NaCl, 3 mM β-mercaptoethanol, 0.1 mM PMSF) followed by centrifugation at 15,000 g for 30 min. Cleared lysate was loaded on 5 mL Ni-NTA column (Qiagen) equilibrated with buffer C, washed with 5 column volumes of buffer C and with 5 column volumes of buffer C containing 20 mM Imidazole. Then, elution with buffer C containing 200 mM Imidazole was carried out. Fractions containing AR9 nvRNAP core or holoenzyme were pooled and diluted ten times by buffer D (20 mM Tris pH 8, 0.5 mM EDTA, 1 mM DTT) or by buffer E (20 mM Bis-tris propane pH 6.8, 0.5 mM EDTA, 1 mM DTT) correspondingly and applied to a MonoQ 10/100 column (GE Healthcare). Bound proteins were eluted with a linear 0.25–0.45 M NaCl gradient in buffer D or E correspondingly.

Cells containing over-expressed recombinant tagless AR9 nvRNAP core were harvested by centrifugation and disrupted by sonication in buffer B followed by centrifugation at 15,000g for 30 min. An 8% polyethyleneimine (PEI) solution (pH 8.0) was added with stirring to the cleared lysate to the final concentration of 0.8%. The resulting suspension was incubated on ice for 1 hour and centrifuged at 10,000g for 15 min. The supernatant was removed and the pellet was resuspended in buffer B containing 0.3 M NaCl. After 10 min incubation, the PEI pellet was formed by centrifugation as previously. Then, supernatant was removed and the pellet was resuspended in buffer B containing 1 M NaCl followed by centrifugation at 10,000g for 15 min. Eluted proteins were precipitated the supernatant by addition of ammonium sulfate to 67% saturation and dissolved in buffer D and loaded on equilibrated 5 mL HiTrap Heparin HP sepharose column (GE Healthcare). The protein was eluted with a linear 0-1 M NaCl gradient in buffer D. Fractions containing tagless AR9 nvRNAP core were pooled and subjected to anion exchange chromatography as described above for AR9 nvRNAP core.

The AR9 nvRNAP core sample was polished and buffer-exchanged using size exclusion chromatography on a Superdex 200 10/300 (GE Healthcare) column equilibrated with buffer D containing 100 mM NaCl. The tagless AR9 nvRNAP core and AR9 nvRNAP holoenzyme were not subjected to size exclusion chromatography – salt concentration in the sample was lowered during the concentration procedure.

The fractions containing AR9 nvRNAP core, tagless AR9 nvRNAP core or holoenzyme were concentrated to a final concentration of 20 mg/mL and used for crystallization or cryo-EM.

### Preparation of the promoter complex for structure determination

To prepare the DNA template for crystallization and cryo-EM, two corresponding oligonucleotides (**Supplementary Table 1**) that were synthetized by IDT with dual PAGE and HPLC purification at a final concentrations of 100 μM each were annealed together by mixing in a buffer containing 20 mM Bis-tris propane pH 6.8, 100 mM NaCl, 4 mM MgCl_2_, 0.5 mM EDTA and incubating at 65 °C for 1 minute and cooling down to 4 °C by a decrement of 1°C per minute. A 1.5-fold molar excess of the DNA template was added to the holoenzyme and incubated for 30 min at room temperature (the final concentrations: 10 mg/mL of the protein (34 μM) and 50 μM of the DNA). The obtained complex was used for crystallization and cryo-EM directly.

### Crystallization of AR9 nvRNAP

The initial crystallization screening was carried out by the sitting drop method in 96 well ARI Intelliwell-2 LR plates using Jena Bioscience crystallization screens at 19°C. PHOENIX pipetting robot (Art Robbins Instruments, USA) was employed for preparing crystallization plates and setting up drops, each containing 200 nL of the protein and the same volume of well solution. Optimization of crystallization conditions was performed in 24 well VDX plates and thin siliconized cover slides (both from Hampton Research) by hanging drop vapor diffusion. The best crystals were obtained as follows: i) an 1.5 μl aliquot of AR9 nvRNAP core (4.5 mg/mL) was mixed with an equal volume of a solution containing 100 mM Tricine pH 8.8, 270 mM KNO_3_, 15 % PEG 6000, 5 mM MgCl_2_, and incubated as a hanging drop over the same solution; ii) an 1.5 μl aliquot of tagless AR9 nvRNAP core (7.5 mg/mL) was mixed with an equal same volume of a solution containing 150 mM Malic acid pH 7, 150 mM NaCl, 14 % PEG 3350 and incubated as a hanging drop over the same solution; iii) an 1.5 μl aliquot of the AR9 nvRNAP promoter complex (10 mg/mL) was mixed with an equal same volume of a solution containing 150 mM MIB pH 5, 150 mM LiCl, 13 % PEG 1500 and incubated as a hanging drop over the same solution. Some crystal reached their final size the next day and some grew for two weeks at 19 C° temperature.

### Preparation of heavy-atom derivative crystals

The following compounds were tested for heavy atom derivatization of AR9 nvRNAP core crystals (by co-crystallization and soaking): SrCl_2_, GdCl_3_, Na_2_WO_4_, HgCl_2_, Pb(NO_3_)_2_, thimerosal (2-(C_2_H_5_HgS)C_6_H_4_CO_2_Na), 10 compounds containing Eu and Yb atoms (JBS Lanthanide Phasing Kit), three compounds containing W (JBS Tungstate Cluster Kit) and one cluster compound containing Ta (Ta_6_Br_12_ JBS Tantalum Cluster Derivatization Kit). The crystals were soaked in a range of concentrations of heavy atom compounds (between 0.1 mM and 100 mM) that were added to the crystallization solution. The soaking time was varied from 2 hours to 2 days. Among all examined conditions, only solutions containing 10 mM thimerosal or 1 mM tantalum bromide resulted in heavy atom derivatization (judging by the presence of anomalous signal in X-ray diffraction data) upon overnight soaking.

To produce a Se-methionine (SeMet) derivative of the AR9 nvRNAP core enzyme, the corresponding plasmid was transformed into B834(DE3) chemically competent *E. coli* cells. The cells were first grown in LB medium until the optical density OD_600_ reached a value of 0.35. The cells were then pelleted by centrifugation at 4,000 g for 10 min at 4°C and transferred to the SelenoMet Medium (Molecular Dimensions) that was supplemented with ampicillin at a concentration of 100 μg/mL. The protein expression then proceeded according to the manufacturer’s instructions. All the subsequent steps were the same as for the native protein.

### X-ray data collection

Cryoprotectant solutions were prepared by replacing 25% of water in the crystallization solution (hanging drop well solution) with ethylene glycol, which was found to be the best cryoprotectant by trial and error. The crystals were either soaked for 1-5 minutes in the cryoprotectant solution or briefly dipped into it and then flash frozen in liquid nitrogen. Such frozen crystals were then transferred to a shipping Dewar and shipped to the APS (LS-CAT) or ALS (BCSB) synchrotrons for remote data collection. X-ray diffraction data and fluorescent spectra were collected in a nitrogen stream at 100 K.

Heavy atom and SeMet derivative data were collected at the absorption peak wavelength (the white line, if present) of the X-ray fluorescence spectrum.

### X-ray structure determination

The structure determination process spanned nearly three years. Initially, we aimed to solve the structure of the AR9 nvRNAP core enzyme by X-ray crystallography or cryo-EM and use it to solve the structure of the promoter complex. However, the atomic model of the promoter complex obtained by cryo-EM was built first. Then, it was used to solve the X-ray structure of the core and to interpret the cryo-EM map of the holoenzyme. The path to the atomic model described below is the trunk of a tree that had many branches representing things that did not work. Many different tools were used in the determination of this structure albeit most of them ended up being dead end branches. The following procedure involves fewest datasets and fewest steps that lead to an interpretable map.

None of the RNAP structures present in the PDB at the start of this project were sufficiently similar to solve the structure of the AR9 nvRNAP core by molecular replacement (MR), so crystallographic phases had to be obtained by a *de novo* phasing procedure (heavy atom isomorphous replacement or anomalous scattering). Severe anisotropy and inconsistent diffraction of AR9 nvRNAP core native crystals made this task extremely complicated and we had to screen hundreds of heavy atom-soaked crystals for diffraction. The SeMet derivative diffracted to 5.5 Å resolution, which was insufficient to solve the Se substructure using anomalous scattering. Moreover, this derivative was not isomorphous to any of the native datasets.

An interpretable map was obtained by a convoluted procedure. A map in which the characteristic features of a DNA-dependent RNAP – two adjoining double-ψ β-barrel (DPBB) domains and several large α-helices, including the bridge helix (although split in the middle) – could be discerned, but no side chain densities were present, was obtained by a multiple isomorphous replacement plus anomalous scattering method which was applied to the native, Ta_6_Br_12_, and thimerosal derivative datasets of the His-tagged AR9 nvRNAP core enzyme (see **Supplementary Table 2**). This map was calculated by the SHARP software package that was run with mostly default settings ^35^. The most similar part^36^ of the archaeal RNAP structure^37^ (PDB code 4ayb) was fitted into this density using Coot^38^ and all parts that did not fit the density and all side chains were removed. The resulting model contained 832 alanine residues.

This model was then used to solve the structure of a unique thimerosal derivative dataset that belonged to a different unit cell (**Supplementary Table 2**) with the help of a MR procedure^39^. This unique dataset resulted from the crystallization of a tagless version of the AR9 nvRNAP core (all native N- and C-termini). It had a very large orthorhombic unit cell with eight molecules of the AR9 nvRNAP core in its asymmetric unit or about 17,800 amino acids. Remarkably, Phaser^40^ was able to locate all eight copies of the AR9 nvRNAP core in this 3.8 Å resolution dataset with help of the 832-residue polyalanine fragment (obtained as described before) as a search model. This polyalanine search model corresponded to about 1.8% of the total protein material in the asymmetric unit and 1% of the total asymmetric unit content if solvent atoms are taken into account. The density was then dramatically improved by 25 cycles of eightfold non-crystallographic symmetry (NCS) averaging using Parrot^41^. The resulting density was mostly continuous and showed many bulky side chains especially in the vicinity of the DPBB domains. Buccaneer^42^ was then used for automatic model building into this map. The Buccaneer model was cleaned up manually and separate chain fragments were assembled into a new intermediate AR9 nvRNAP core model that contained 995 residues of which 937 had side chains. The new intermediate model was then used as a search model in a new round of MR by Phaser that was followed by NCS averaging using Parrot.

The new density was of sufficient quality to recognize the identity of many side chains and for manual model building using Coot. Structures of the *Mycobacterium tuberculosis* and *E. coli* RNAPs (PDB codes 5ZX3^43^ and 6C9Y^44^, respectively) were used to aid in chain tracing. Additionally, this thimerosal derivative dataset contained Hg atoms, identified with the help of anomalous Fourier synthesis, that were expected to bind to cysteine side chains. Thus, the Hg atoms were used as markers to maintain the chain register.

Eventually, a large fraction of the nvRNAP core atomic model was complete. Some peripheral parts, however, and the β′C gp154 Insertion domain were too disordered for model building. While this work was in progress, an interpretable (3.8 Å resolution) cryo-EM map of the AR9 nvRNAP promoter complex was obtained. This map was of better quality than the 3.8 Å resolution, large unit cell, X-ray dataset of the tagless core enzyme, so further rounds of model building were continued using the cryo-EM map. Once the holoenzyme model was complete, the core or the entire holoenzyme (but not the DNA) were used to solve the crystal structure of the native AR9 nvRNAP core (3.3 Å resolution, the standard unit cell, **Supplementary Table 2**) and promoter complex (3.4 Å resolution, **Supplementary Table 2**) by MR using Phaser.

Some peripheral regions of the cryo-EM promoter complex map, namely, the β′C gp154 Insertion domain, residues 130-289 of βN gp105, and the peripheral parts of gp226, were too poor for reliable *de novo* model building. Fortunately, the structure of AR9 nvRNAP promoter complex was one of large multisubunit targets of the CASP14 protein structure prediction competition. The Google DeepMind AlphaFold2 software predicted the structure of difficult-to-build domains with excellent accuracy^23^. This allowed us to complete the AR9 nvRNAP promoter complex model in the cryoEM map first and then use this model to solve the 3.4 Å resolution crystal structure of the promoter complex.

Refinement of crystallographic and cryo-EM models was performed using Phenix^45^ and Coot. Additional details describing model building and the analysis of AlphaFold2 models are given elsewhere^46^.

### Cryo-EM sample preparation and data acquisition of the AR9 nvRNAP promoter complex

QUANTIFOIL 1.2/1.3 copper grids were plasma cleaned for 30s using the model 950 advanced plasma system by Gatan. 3 μL of 10 mg/mL of the AR9 nvRNAP holoenzyme (34 μM) and 50 μM of the DNA nucleotide in 20 mM Bis-tris propane pH 6.8, 100 mM NaCl, 2 mM MgCl_2_, 0.5 mM EDTA was pipetted onto to the grid and blotted using a Vitrobot (Thermo Fischer Scientific) at 100% humidity for 5 s. Following blotting, the sample was plunged into liquid ethane cooled by liquid nitrogen.

5,351 (5,760 × 4,092 pixels) micrograph movies were collected using the EPU software on a Titan Krios 300kV electron microscope with a BioQuantum K3 imaging filter with a 20-eV slit. Each movie contained 56 frames collected over 1.5 s, with a frame dose of 0.78 e/Å^2^ and pixel size of 1.09 Å. Movies were collected over a defocus range of −1 to −4 μm.

### Cryo-EM image processing of the AR9 nvRNAP promoter complex

Image processing was performed using Eman2 ^47^, Relion3.0 ^48^ and CryoSPARC3.0 ^49^ (**Supplementary Fig. 3, Supplementary Table 3**). All movies were motion corrected using MotionCor2 ^50^. Estimation of the contrast transfer function (CTF) parameters was performed by CTFFIND4.1 ^51^ over the resolution range of 5.0-30.0 Å. E2boxer ^47^ was used for particle picking, resulting in 420,791 particles with a box size of 300 pixels. Particle box coordinates were used by Relion3.0 to extract the boxes. 2D classification by Relion3.0 resulted in 243,976 particles belonging to high quality classes. These particles were imported into CryoSPARC3.0, where *ab initio* reconstruction was carried out using three models. Following the *ab initio* reconstruction, 3D classification was performed with five classes, a box size of 150 pixels to improve speed, a batch size of 2,000 particles per class and an assignment convergence criterion of 2%. Non-Uniform (NU) refinement ^52^ was executed using both the 106,867 particles and the map from the most populous class of 3D classification. 3D local refinement was then carried out using an alignment resolution of 0.25° and NU refinement. Subsequently, local CTF-refinement was performed using a search range of 3.5-20.0 Å. Following this, around round of NU refinement and 3D local refinement with an alignment resolution of 0.25° and NU-refinement was performed. The resulting map was sharpened with a B-factor of −138 Å^2^.

### Cryo-EM sample preparation and data acquisition of the AR9 nvRNAP holoenzyme

TED PELLA 200 mesh PELCO NetMesh copper grids were plasma cleaned for 30s using the model 950 advanced plasma system by Gatan. 3 μl of 20mg/ml His-tag 5s nvRNAP in 20 mM Bis-tris propane pH 6.8, 100 mM NaCl, 4 mM MgCl_2_, 0.5 mM EDTA buffer was pipetted onto to the grid and blotted using a Vitrobot (Thermo Fischer Scientific) at 100% humidity for 5 s. Following blotting, the sample was plunged into liquid ethane cooled by liquid nitrogen.

2,691 (3,838 × 3,710 pixels) micrograph movies were collected using the EPU software on a Titan Krios 300kV electron microscope with a K2 Summit camera and a 20-eV slit. Each movie contained 40 frames collected over 8 s, with a frame dose of 1.08 e/Å^2^ and pixel size of 1.08 Å. Movies were collected with a target defocus of −1.8 μm. Images were collected with a 30-degree tilt.

### Cryo-EM image processing of the AR9 nvRNAP holoenzyme

Image processing was performed using Relion2.0 ^48^. All movies were motion corrected using MotionCor2 ^50^. Estimation of the contrast transfer function (CTF) parameters was performed by Gctf^53^. Particle picking resulted in 227,577 particles with a box size of 200 pixels. 2D and 3D classification by Relion2.0 resulted in 104,471 particles belonging to high quality classes. The resulting particles were refined to a resolution of 4.4 Å. The resulting map was sharpened with a B-factor of −87 Å^2^.

### Gp226 cloning, purification and limited digestion with trypsin

The AR9 gene *226* was PCR amplified from AR9 genomic DNA and cloned into the pQE-2 vector (QIAGEN) between the SacI and SalI restriction sites. The resulting plasmid was transformed into BL21 (DE3) chemically competent *E. coli* cells. The culture (7 L) was grown at 37°C to an OD_600_ of 0.5 in LB medium supplemented with ampicillin at a concentration of 100 μg/mL, and recombinant protein overexpression was induced with 1 mM IPTG for 4 hours at 22°C. Cells containing over-expressed recombinant protein were harvested by centrifugation and disrupted by sonication in buffer C followed by centrifugation at 15,000 g for 30 min. Cleared lysate was loaded on a 5 mL Ni-NTA column (Qiagen) equilibrated with buffer C, washed with 5 column volumes of buffer C and with 5 column volumes of buffer C containing 20 mM Imidazole. Then, elution with buffer C containing 200 mM Imidazole was carried out. Fractions containing gp226 were pooled, concentrated and subjected to gel-filtration on a Superdex 200 10/300 (GE Healthcare) column equilibrated with buffer С. The fractions containing gp226 monomer were concentrated to a final concentration of 1 mg/mL and used for the limited proteolysis experiment.

Trypsin digestion of gp226, which had a concentration of 80 ng/μL, was carried out in 20 ul of the digestion buffer (50 mM NaH_2_PO_4_ pH 8.0, 300 mM NaCl) that contained a range of trypsin concentrations (Sigma-Aldrich). The trypsin to gp226 molar ratios were from 0.03 to 0.6. The reactions were allowed to proceed for 1 hour at 25°C and stopped by the addition of Laemmli loading buffer and immediate boiling. The reaction products were analyzed by denaturing SDS polyacrylamide gel electrophoresis (SDS-PAGE) with subsequent mass-spectrometry as described previously^1^.

### DNA templates for transcription assay

Long DNA templates containing late AR9 promoters were prepared by polymerase chain reaction (PCR). PCRs were done with Encyclo DNA polymerase (Evrogen, Moscow) and the AR9 genomic DNA as a template, with a standard concentration of dNTPs (Thermo Fisher Scientific) to obtain DNA fragments with thymine or in the presence of dUTP (Thermo Fisher Scientific) in place of dTTP to obtain DNA fragments with uracil. Oligonucleotide primers used for PCR are listed in (**Supplementary Table 1**).

Short double-stranded and partially single-stranded DNA templates containing the P077 promoter with uracils and thymines at certain positions were prepared by annealing of oligonucleotides ordered from Evrogen (Moscow) and listed in (**Supplementary Table 1**). To prepare specific DNA templates, two corresponding oligonucleotides were annealed together by mixing in buffer containing 40 mM Tris-HCl pH 8, 10 mM MgCl_2_ and 0.5 mM DTT, incubating at 75 °C for 1 minute and cooling down to 4 °C by a decrement of 1°C per minute.

### *In vitro* transcription

Multiple-round run-off transcription reactions were performed in 5 μL of transcription buffer (40 mM Tris-HCl pH 8, 10 mM MgCl_2_, 0.5 mM DTT, 100 μg/mL bovine serum albumin (Thermo Fisher Scientific), and 1 U/μL RiboLock RNase Inhibitor (Thermo Fisher Scientific)) and contained 100 nM AR9 nvRNAP holoenzyme and 100 nM DNA template. The reactions were incubated for 10 min at 37°C, followed by the addition of 100 μM each of ATP, CTP, and GTP, 10 μM UTP and 3 μCi [α-^32^P]UTP (3000 Ci/mmol) (**Extended data Fig. 1c, Figure 2b**) or 100 μM each of ATP, UTP, GTP, 10 μM CTP and 3 μCi [α-^32^P]CTP (3000 Ci/mmol) (**Figure 2d, 2f**). Reactions proceeded for 30 min at 37°C and were terminated by the addition of an equal volume of denaturing loading buffer (95% formamide, 18 mM EDTA, 0.25% SDS, 0.025% xylene cyanol, 0.025% bromophenol blue). The reaction products were resolved by electrophoresis on 6-23 % (w/v) polyacrylamide gel containing 8 M urea. The results were visualized with a Typhoon FLA 9500 scanner (GE Healthcare).

### Molecular dynamics general methods

Simulations were carried out on both the LS5 and Stampede2 systems at the Texas Advanced Computing Center (TACC) using NAMD 2.10 ^33^. The CHARMM36 force field was used^54^. Production runs were performed in the isothermal isobaric (NPT) ensemble using Langevin dynamics and a Langevin piston^55^. Alchemical transformations were analyzed by the ParseFEP package^56^. Entropic restraints were calculated via thermodynamic integration. Collective variables were implemented via the colvars module in NAMD^57^. The energy of non-bonded VdW interactions for distances exceeding 10 Å was smoothly decreased to equal zero at 12 Å. A 2 fs timestep was used in all simulations. Long range electrostatics was calculated with the help of the Particle Mesh Ewald algorithm ^58^. During alchemical transformation, a soft core VdW radius of 4 Å^3^ was used to improve convergence and accuracy ^59,60^. Both alchemical and restraint calculation simulations were carried out bidirectionally. The Bennett Acceptance Ratio maximum likelihood estimate^61^ was used to determine free energy change for alchemical transformations. The double decoupling method (DDM) was implemented as described^31,32^.

### Molecular dynamics system setup

The protein structure description files for both structures – the AR9 nvRNAP holoenzyme in complex with the 3′-^−11^UUG^−9^-5′ oligonucleotide bound to the promoter binding pocket and for the 3′-^−11^UUG^−9^-5′ oligonucleotide in the promoter bound conformation – were generated using the psfgen plugin of VMD ^62^. Solvation was performed using TIP3 water^63^ in a box with 15 Å padding in each direction and ionized in 0.1 M NaCl. The holoenzyme-DNA and DNA systems were enclosed in periodic boxes with cell dimensions of (176 Å, 147 Å, 133 Å) and (42 Å, 42 Å, 41 Å), respectively, and contained 91,177 and 2,166 water molecules, respectively. Both systems were first minimized for 1,000,000 steps while restraints and constraints on the protein (in the holoenzyme system), DNA, and water atoms were gradually removed. Both systems were heated from 0 K to 300 K in 5 K increments for a total of 19.2 ns with constraints on backbone atoms. The holoenzyme-DNA and DNA systems were then equilibrated with minimal constraints in the isothermal-isobaric (NPT) ensemble for 100 ns and 50 ns, respectively.

### Definition of collective variables

For the implementation of the DDM method via alchemical transformations, the system must be (harmonically) constrained such that the finite sampling can be focused on relevant regions of phase space. The entropic cost of applying these restraints is evaluated in separate independent simulations. The phase space and the entropic cost are connected to each other through a set of collective variables that are applied to atoms during simulations.

Only one collective variable is needed to restrain the conformation of the oligonucleotide in bulk water (the bulk water DNA system): the root mean squared deviation (RMSD) of all non-H DNA atoms relative to the equilibrated state. In the holoenzyme-DNA complex, seven collective variables are required – six to define the orientation and position of the rigid DNA molecule relative to the holoenzyme complex and one to define the conformation of the DNA. Similarly to the bulk water DNA case, the RMSD of all non-H DNA atoms relative to the equilibrated state is used as the collective variable to restrain the conformation of DNA. We characterize the orientation of the rigid DNA molecule via the relative position of the backbone atoms of ^−11^UUG^−9^ to those of gp226 V206, gp105 N382 and gp226 K262. Accordingly, the orientation of DNA relative to the holoenzyme is characterized by six collective variables: *r* – the distance between ^−11^U and gp226 V206); *ϕ* – the angle between gp226 V206, ^−11^U, and ^−9^G); *χ* – the angle between ^−11^U, gp226 V206, and gp105 N382); *θ* – the dihedral angle between gp226 V206, ^−11^U, ^−9^G, and ^−10^U); *ψ* – the dihedral angle between gp105 N382, gp226 V206, ^−11^U, and ^−9^G); *ξ* – the dihedral angle between ^−11^U, gp226 V206, gp105 N382, and gp226 K262).

The harmonic force constraint constants applied to the distance-type (RMSD and *r*) and angular collective variables were 10 kcal/mol/Å^2^ and 0.1 kcal/mol/deg^2^, respectively. The equilibrium positions for all harmonic restraints were derived from the equilibrated holoenzyme-DNA structure. For restraint estimation simulations, harmonic forces were varied smoothly using a target force exponent value of 4.0. The lambda schedule focused near the value 1.0 to improve simulation convergence and ensure thermodynamic micro-reversibility^64,65^: [1.00, 0.999, 0.99, 0.95, 0.90, 0.85, 0.80, 0.75, 0.70, 0.65, 0.60, 0.55, 0.50, 0.45, 0.40, 0.35, 0.30, 0.20, 0.10, 0.00]. The reverse sequence was used for the backward simulation.

### Thermodynamic cycle

The standard binding free energy was calculated by combining the results of four separate simulations which represent the four vertical reactions of the thermodynamic cycle (**Extended data Fig. 8a**). These simulations evaluate the following parameters (**Supplementary Table 4**): 1) the entropic cost of restraining the promoter DNA to the “bound” state in the promoter binding pocket by adding/removing conformational restraints on the promoter DNA 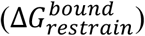; 2) the free energy of coupling/decoupling the promoter DNA from the binding pocket via alchemical transformations with restraints on the conformation of the promoter DNA 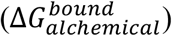; 3) the entropic cost of restraining the promoter DNA to the bound conformation in bulk water by adding/removing conformational restraints on the promoter DNA 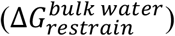; 4) the free energy of coupling/decoupling the promoter DNA from bulk water via alchemical transformations with restraints on the conformation of the promoter DNA 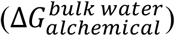. The change in free energy resulting from these transitions is then calculated via thermodynamic integration^66^ and free energy perturbation^67^ methods. The results were validated by checking for micro-reversibility and the absence of hysteresis^56,61^. Following the completion of the thermodynamic cycle, the standard binding free energy of promoter DNA to the holoenzyme binding pocket was found to be −6.9 ± 2.8 kcal/mol.

### Molecular dynamics error analysis

An upper bound on the error of 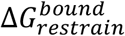, 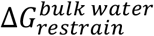 and 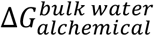 was determined by the hysteresis between backward and forward simulations^32^. The error in 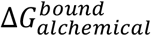 was determined by performing 3 replicates of the simulation and evaluating the standard deviation (**Supplementary Table 4**).

### Data availability

All macromolecular structure data described in this paper have been deposited to the Protein Data Bank and Electron Microscopy Data Bank under the following accession numbers: PDB code 7S00 (AR9 nvRNAP core X-ray structure); PDB code 7S01 (AR9 nvRNAP promoter complex X-ray structure); EMDB code EMD-24765 (AR9 nvRNAP holoenzyme cryo-EM density); EMDB code EMD-24763 (AR9 nvRNAP promoter complex cryo-EM density). Publicly available protein atomic models with the following PDB codes were used in the study: 4AYB^37^, 5ZX3^43^, 6C9Y^44^, 5IPM^15^, 6JBQ^16^, and 7OGP^21^.

## Acknowledgments

This work was supported by Skoltech NGP Program (Skoltech-MIT joint project) and by the Russian Science Foundation (Grant 19-74-00011 to M. L. Sokolova). The work was also supported by the UTMB Department of Biochemistry and Molecular Biology and by the UTMB Sealy Center for Structural Biology and Molecular Biophysics. The MD work was performed using the computing facilities of the Texas Advanced Computing Center (TACC, http://www.tacc.utexas.edu) at The University of Texas for which we are very grateful. We thank the Stanford-SLAC Cryo-EM Facilities, supported by Stanford University, SLAC and the National Institutes of Health S10 Instrumentation Programs that were used to collect the AR9 nvRNAP holoenzyme cryo-EM data. We acknowledge the use of the Advanced Photon Source, a U.S. Department of Energy (DOE) Office of Science User Facility operated for the DOE Office of Science by Argonne National Laboratory under Contract No. DE-AC02-06CH11357. We thank the staff of the LS-CAT Sector 21 beamlines that is supported by the Michigan Economic Development Corporation and the Michigan Technology Tri-Corridor (Grant 085P1000817). We acknowledge the use of the Berkeley Center for Structural Biology (supported in part by the Howard Hughes Medical Institute) at the Advanced Light Source (a Department of Energy Office of Science User Facility under Contract No. DE-AC02-05CH11231) and we thank the staff of the beamline 5.0.2. Finally, we thank Dr. Mark A. White for his help and assistance with the initial crystallization and X-ray data collection of the AR9 nvRNAP core and Dr. Michael B. Sherman for his help with the cryo-EM data collection of all datasets used in this paper. The research reported in this paper extensively used the facilities and resources of the UTMB SCSB Macromolecular Structure X-ray Laboratory and UTMB SCSB Cryo-EM Laboratory.

## Author contributions

**K.V.S.** and **M.L.S.** conceived the study. **M.L.S.** cloned, purified and crystallized AR9 nvRNAP core, tagless AR9 nvRNAP core, and AR9 nvRNAP holoenzyme in complex with promoter DNA, derivatized crystals, prepared samples for cryo-EM, purified gp226 and performed limited trypsinolisis. **A.F**. obtained and analyzed all cryo-EM reconstructions, built parts of atomic models, and performed all MD work. **A.V.D.** purified AR9 nvRNAP holoenzyme and its mutants and performed *in vitro* transcription assays under the supervision of **M.L.S**. **J.G.** under the supervision of **M.L.S.** crystallized the AR9 nvRNAP holoenzyme in complex with promoter DNA. **T.O.** performed mass-spectrometry. **P.G.L.** collected X-ray data, solved all crystal structures, and built and refined all atomic models. The **AF team** created models of all five AR9 nvRNAP holoenzyme proteins that were used by **P.G.L.** and **A.F.** in the interpretation of cryo-EM and X-ray crystallography electron density maps. **A.F., M.L.S.**, **S.B.**, and **P.G.L.** analyzed the structures. **P.G.L.** and **A.F.** wrote the manuscript, which was read, edited, and approved by all authors.

## Supplementary Information

is available for this paper.

## Correspondence and requests

for materials should be addressed to Maria L. Sokolova, Petr G. Leiman, or Konstantin V. Severinov.

## Extended Data Figure Legends

**Extended Data Fig. 1.**
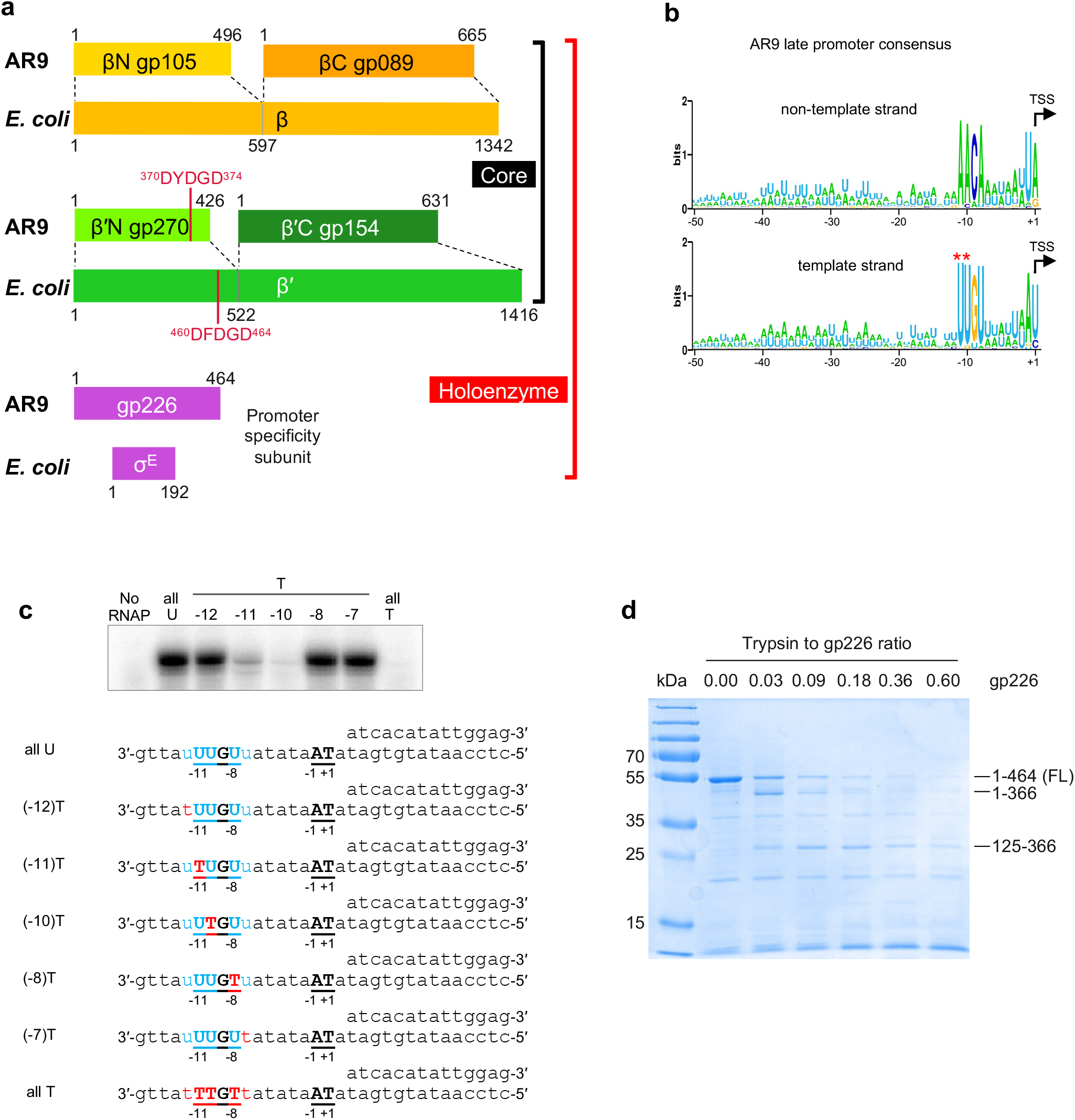
Organization and promoter consensus of the AR9 nvRNAP. **a**, Organization of the catalytically active core and promoter initiation competent holoenzyme of the AR9 nvRNAP. A pair of genes encode a protein complex that is homologous to the bacterial subunits β or β′. The promoter specificity subunit displays no detectable sequence similarity to bacterial sigma factors. **b**, The consensus of AR9 late promoters recognized by the nvRNAP. Both DNA strands are shown. **c**, The dependance of the AR9 nvRNAP *in vitro* transcription activity on the position and number of T bases in the promoter, which is located in the template strand of DNA (bold-underlined). **d**, The resistance of recombinantly expressed gp226 to proteolysis by trypsin. The identities and sizes of labeled major products, given as residue ranges, have been established using mass spectrometry. FL stands for the full length protein. See also **Supplementary Fig. 1**. Two technical replicates of two biological replicates of the *in vitro* transcription and trypsin proteolysis experiments resulted in similar outcomes and one of them is shown. The uncropped autoradiogram and SDS-PAGE are presented in **Supplementary Fig. 1**.

**Extended Data Fig. 2.**
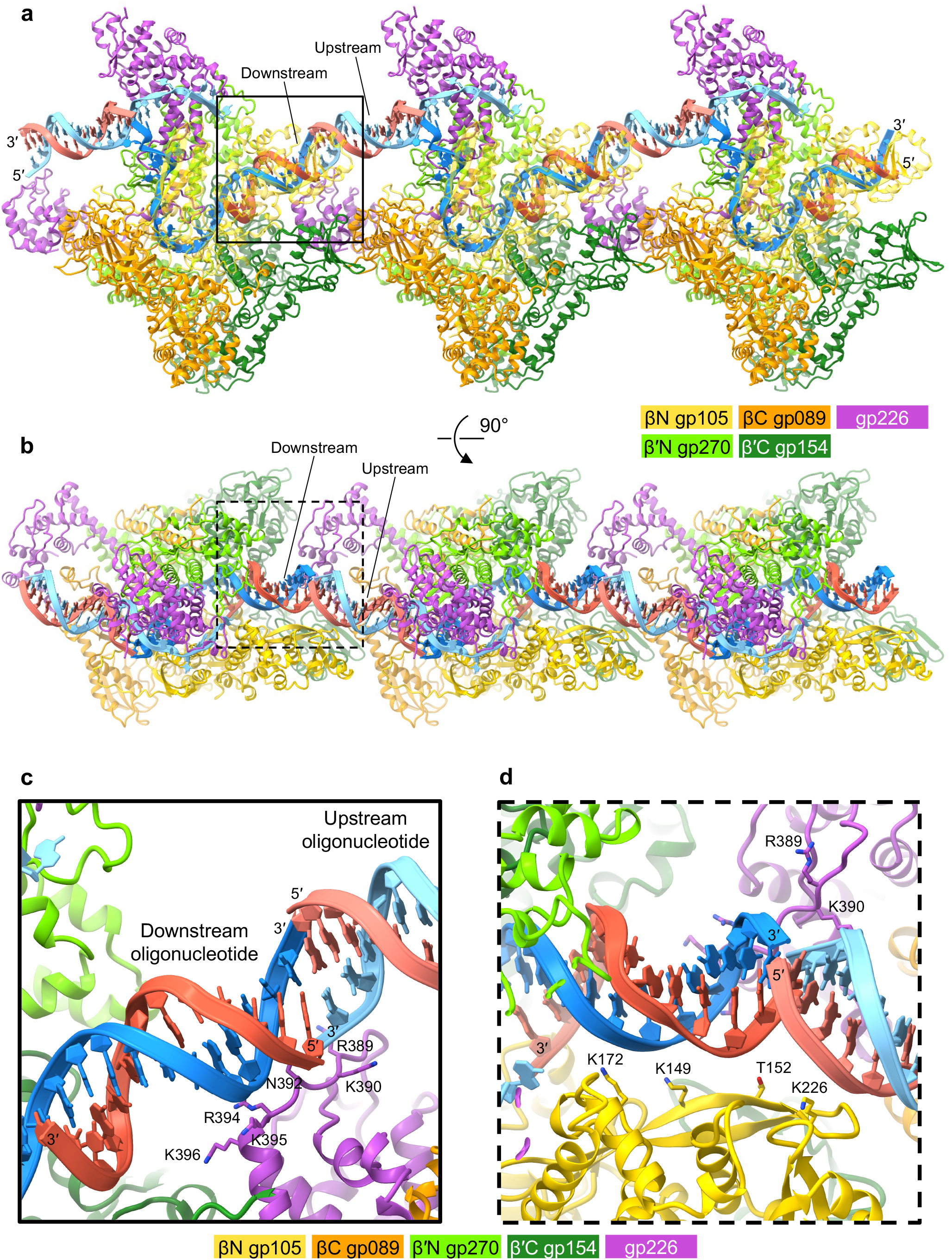
AR9 nvRNAPs and up- and downstream oligonucleotides form a train *in crystallo*. **a** and **b**, Two orthogonal views of three nvRNAPs demonstrating the peculiar crystal packing. **c** and **d**, Two areas of interest indicated with a solid and dashed line square in panels **a** and **b** that show details of pi-pi stacking interactions between the ends of the up- and downstream oligonucleotides. Residues most proximal to the DNA are shown in a stick representation and labeled. In both panels, the color code is as in **Fig. 1a**.

**Extended Data Fig. 3.**
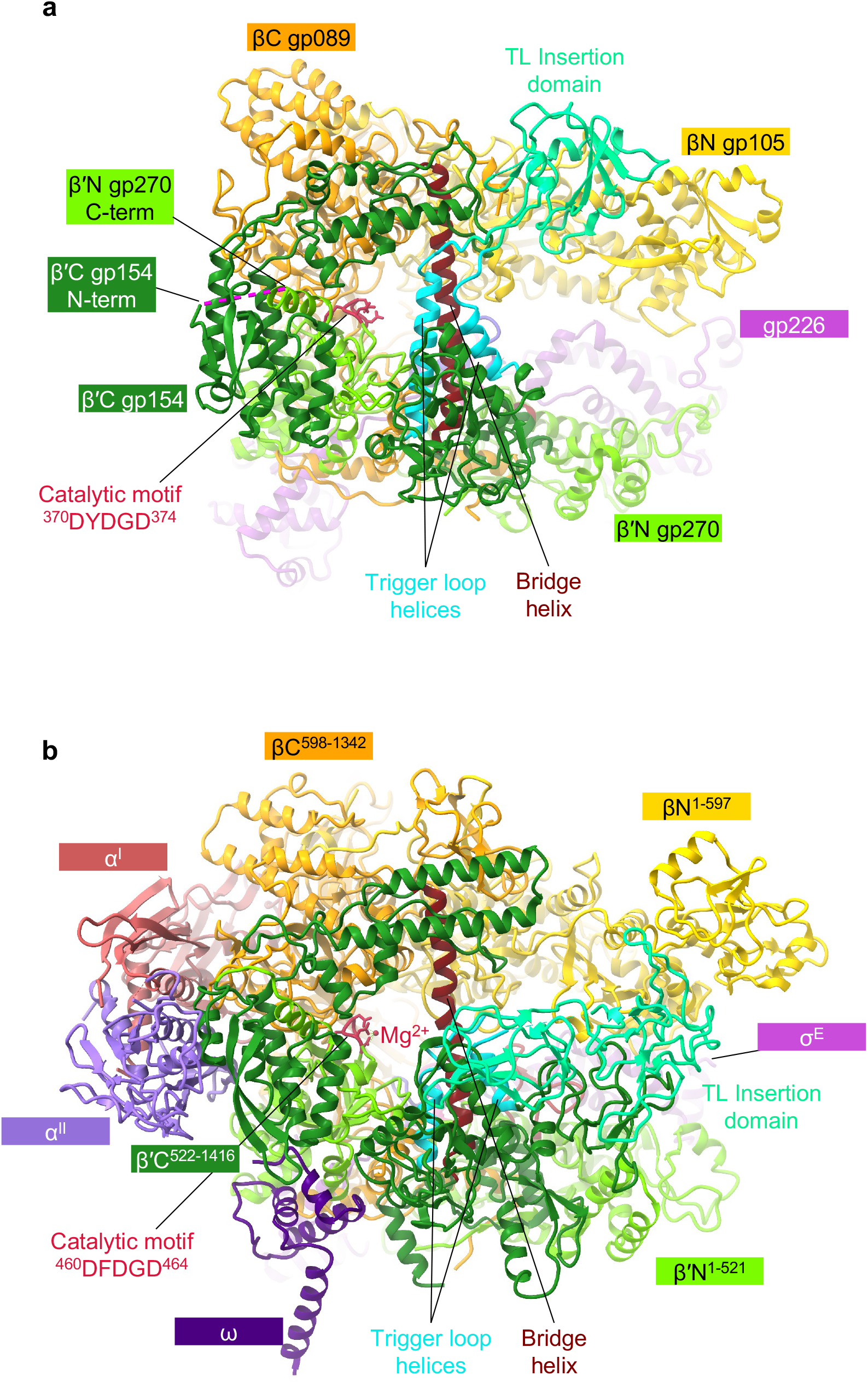
Comparison of the AR9 nvRNAP and *E. coli* RNAP-σ^E^ holoenzymes. **a** and **b**, Ribbon diagrams of the AR9 nvRNAP and RNAP-σ^E^ (PDB code 6JBQ^16^), respectively, are viewed from the NTP entrance channel. Nucleic acids are not shown for clarity. Gp226 and σ^E^ extend into the plane of the paper and are almost completely obscured by the depth-cueing effect. In both molecules, key elements are colored similarly and labeled. In panel **a**, the color code is as in **Fig. 1a**.

**Extended Data Fig. 4.**
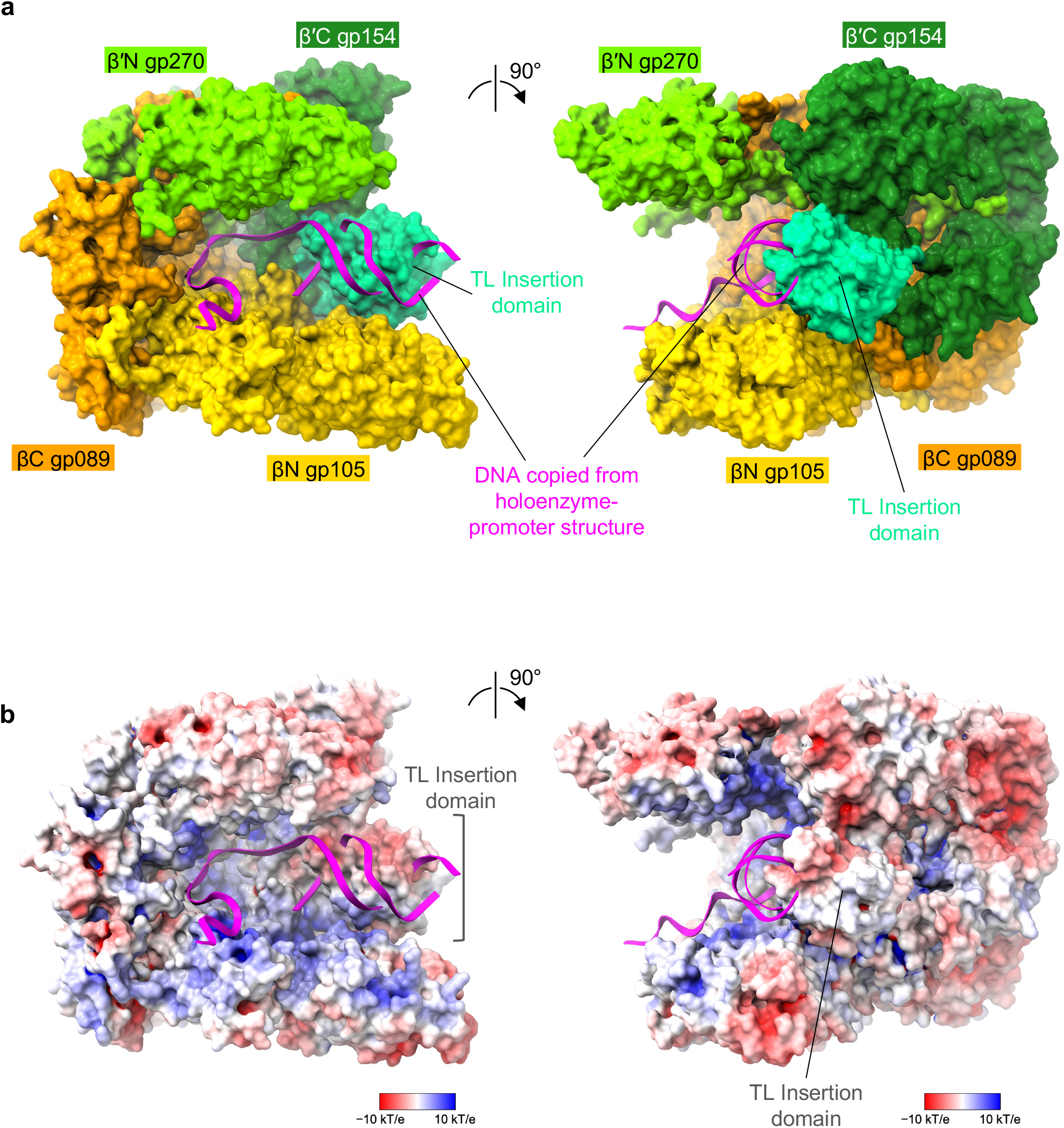
Structure of the AR9 nvRNAP core. **a**, Molecular surface of the AR9 nvRNAP core crystal structure, with subunits colored as in **Fig. 1a**, with the downstream DNA oligonucleotide copied from the holoenzyme-promoter structure. The insertion domain partially obstructs the DNA binding cleft. **b**, Electrostatic potential is mapped onto the molecular surface of the AR9 nvRNAP core. The orientations are identical to panels **a**.

**Extended Data Fig. 5.**
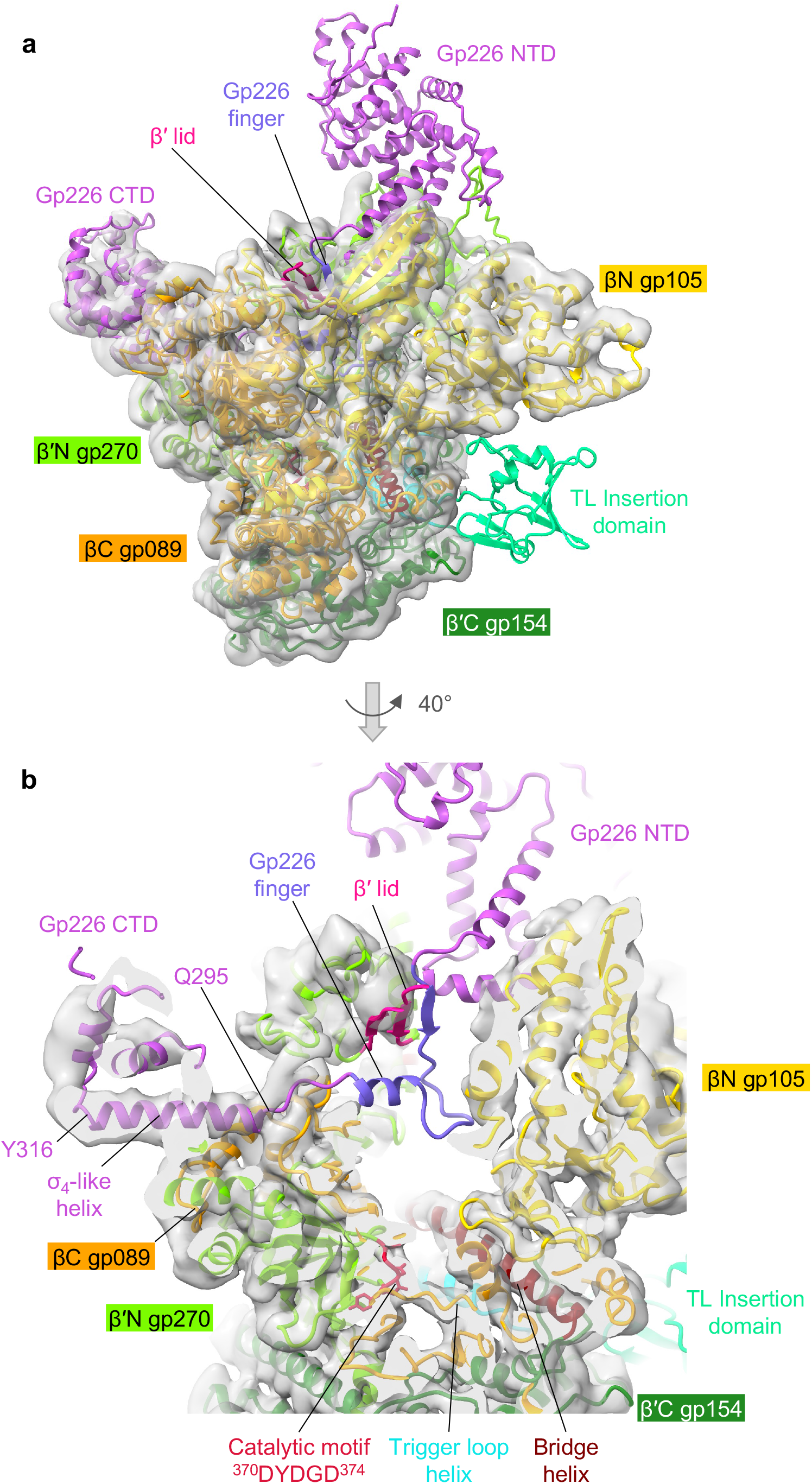
Cryo-EM structure of the AR9 nvRNAP holoenzyme. **a**, Cryo-EM map of the AR9 nvRNAP holoenzyme contoured at 4.0 std dev above the mean (semitransparent gray) with the fitted atomic model of the promoter complex (sans DNA) colored as in **Fig. 1a**. **b**, A zoomed-in view of the catalytic cleft demonstrating the degree of gp226 disorder.

**Extended Data Fig. 6.**
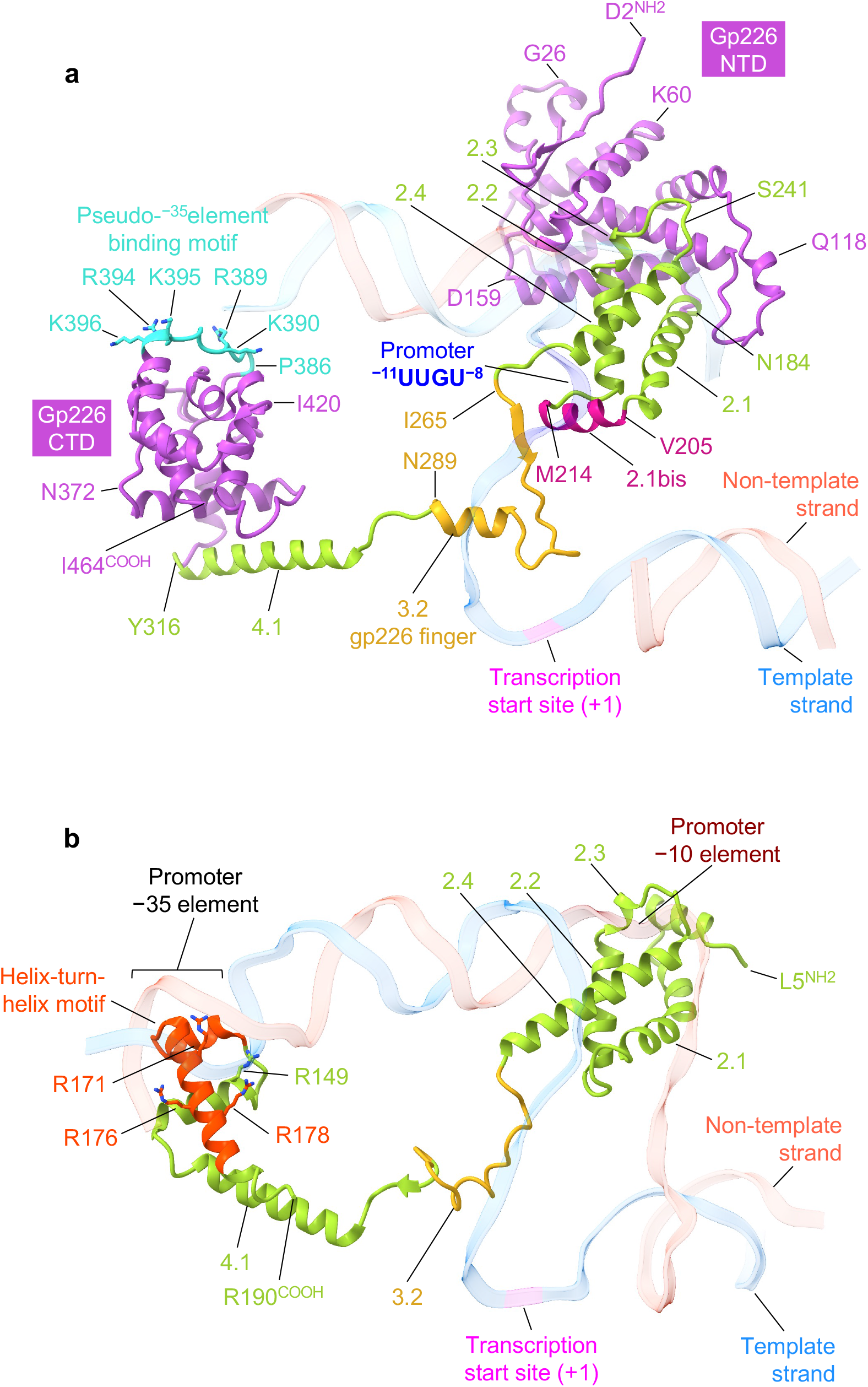
Structure of the promoter specificity subunit gp226. **a**, Ribbon diagram of gp226 with regions structurally similar to bacterial σ factors colored in yellow green (helices 2.1 through 2.4 and 4.1) and gold (finger). The unique N- and C-terminal domains colored medium orchid. Residue numbers and identities are given at key locations. The DNA strands are colored as in **Fig. 1a** and **1b** and are semitransparent. The pseudo^−35^-element binding motif is turquoise and its positively charged and solvent exposed residues are shown in a stick representation. **b**, Ribbon diagram of the *E. coli* σ^E^ factor (PDB code 6JBQ^16^) with its helix-turn-helix motif colored in orange red. Positively charged residues that interact with the DNA are shown in a stick representation and labeled. The DNA backbone is semitransparent.

**Extended Data Fig. 7.**
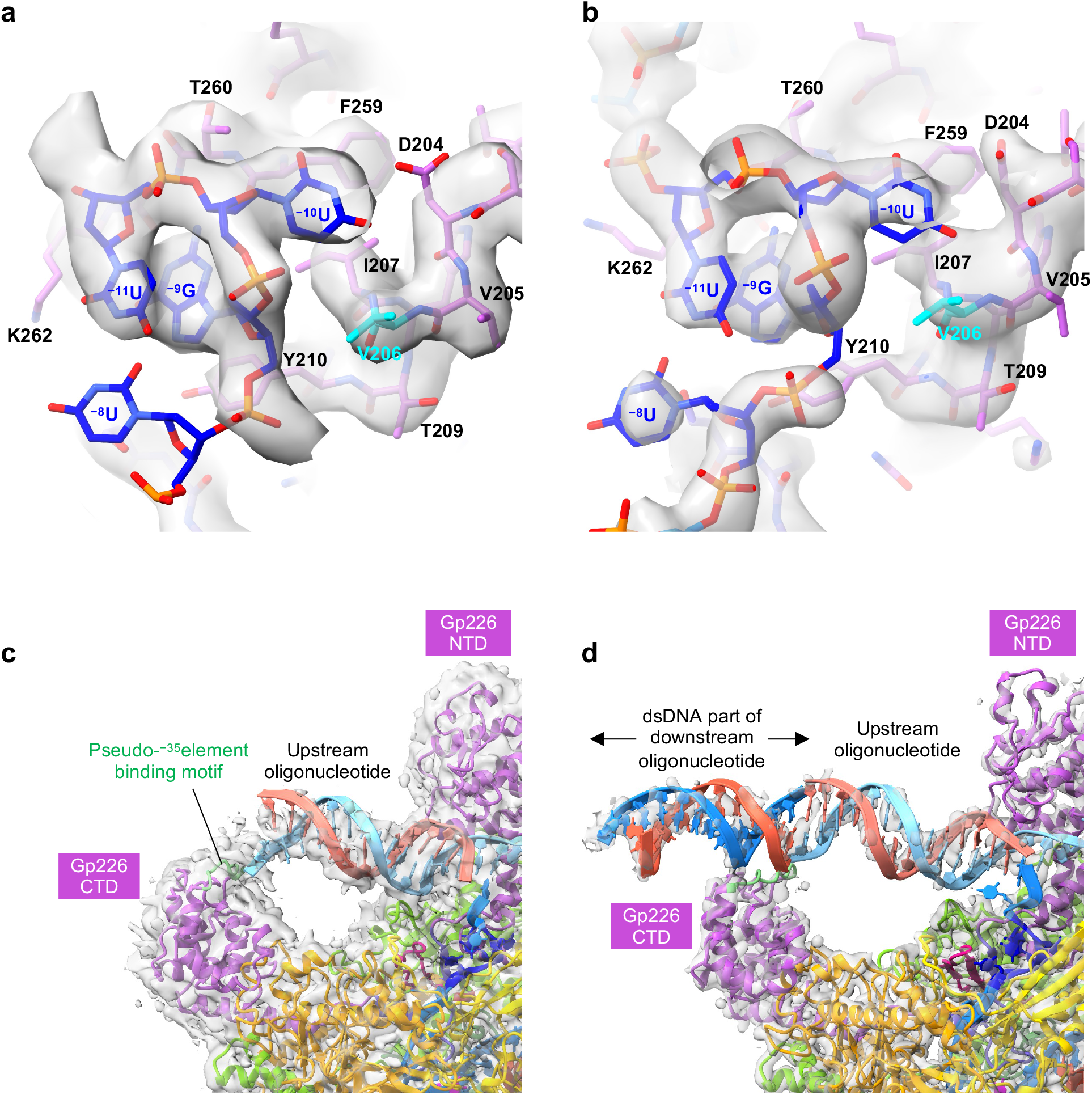
Electron density of the AR9 nvRNAP-DNA interacting regions. **a** and **b**, Cryo-EM and composite omit X-ray electron density maps of the promoter binding pocket with refined atomic models, respectively. The cryo-EM and X-ray maps are contoured at 5.0 and 1.5 std dev above the mean, respectively. The carbon atoms are colored as in **Fig. 1a**. V206, which is critical for U specificity, is colored cyan. **c** and **d**, Cryo-EM and composite omit X-ray electron density maps of the gp226 CTD with refined atomic models, respectively. The cryo-EM and X-ray maps are contoured at 2.0 and 1.0 std dev above the mean, respectively. The ds segment of the downstream oligonucleotide belonging to a neighboring unit cell is shown in the X-ray map (see **Extended data Fig. 2**). Proteins are colored as in **Fig. 1a**. The pseudo-^−35^element binding motif is colored light green.

**Extended Data Fig. 8.**
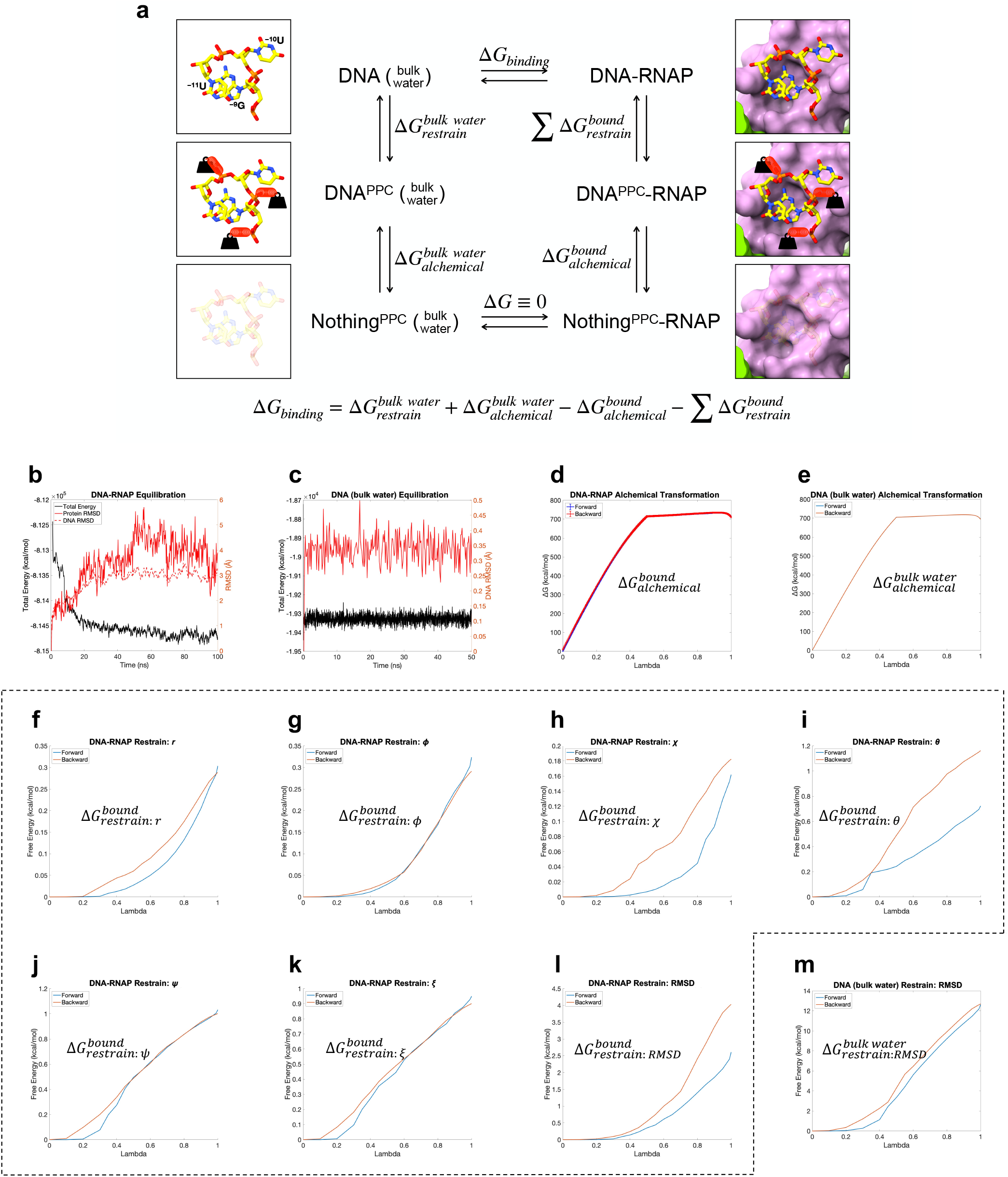
Derivation of the promoter binding free energy using molecular dynamics. **a,** Thermodynamic cycle of promoter binding. The PPC superscript (e.g. DNA^PPC^) stands for Promoter Pocket Conformation in regard to the structure of the 3′-^−11^UUG^−9^-5′ DNA trinucleotide. **b,** Equilibration and relaxation of the cryo-EM derived atomic model of the AR9 nvRNAP holoenzyme with the 3′-^−11^UUG^−9^-5′ DNA trinucleotide bound to the promoter pocket. **c**, Equilibration and relaxation of the 3′-^−11^UUG^−9^-5′ trinucleotide in bulk water. **d** and **e**, Energetics of forward and backward alchemical transformations of the 3′-^−11^UUG^−9^-5′ DNA trinucleotide in the promoter pocket of the AR9 nvRNAP holoenzyme and in the PPC in bulk water, respectively. **f, g, h, i, j, k, l**, Entropic cost of applying seven harmonic constraints to the 3′-^−11^UUG^−9^-5′ DNA trinucleotide to maintain it in the promoter pocket-bound state. **m**, Entropic cost of the harmonic RMSD constraint on the 3′-^−11^UUG^−9^-5′ DNA trinucleotide to maintain the PPC.

**Extended Data Fig. 9.**
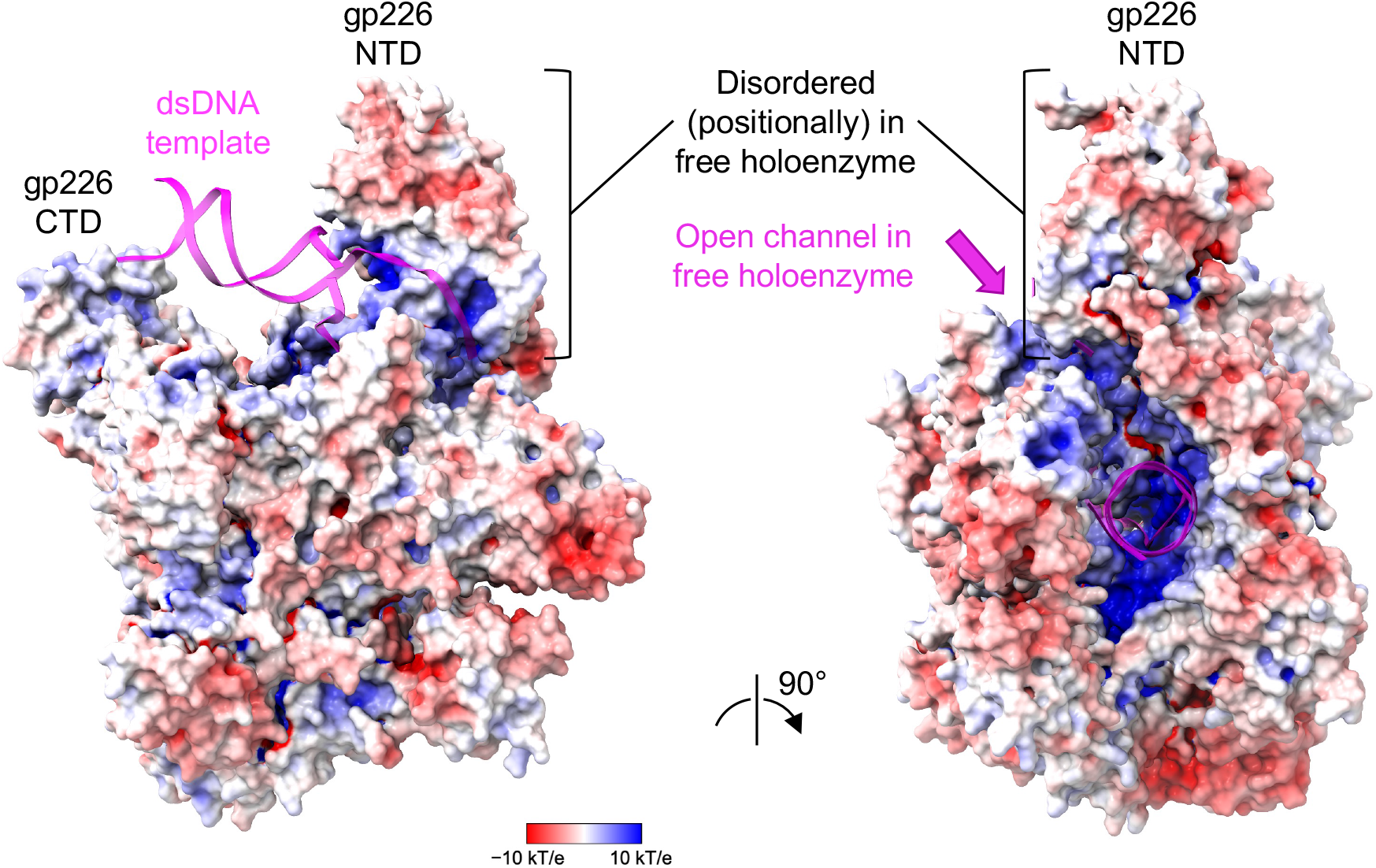
Distribution of electrostatic potential on the surface of the AR9 nvRNAP promoter complex. The DNA (colored magenta) was excluded from the calculations. The orientation of the molecule is as in **Fig. 1a**.

