## Supplementary Information for "Template strand deoxyuridine promoter recognition by a viral RNA polymerase"

### Contents

**Supplementary Tables 1-4**

**Supplementary Figures 1-3**

### Supplementary Table 1. Oligonucleotides used in the study.

| **PCR primers used for site-directed mutagenesis of gp226 (Figure 2b, 2d, 2f)** | | |
|---|---|---|
| *Mutation* | *Oligonucleotide name* | *5′-3′ sequence* |
| gp226 V206G | gp226-V206X-rev | aacatctgagtactttgtctcgaagacgcga |
| | gp226-V206G-dir | gagacaaagtactcagatgttggaatctggacgtaccttaaaaac |
| $A^5$ mutant (gp226 R389A, K390A, R394A, K395A, K396A) | gp226-R389X-rev | cacatttggtgcgatagcgcaggataaaatctc |
| | gp226-A-for | gctatcgcaccaaatgtggccgcgatgaacaacgctgcagcattagttgataaaatcatc |
| gp226 Y246A | gp226_Y246A_F | atctccgcgcttgacgttgtgatcaaacagaaattg |
| | gp226_Y246A_R | gtcaagcgcggagattaccgagctattgtgtttc |
| gp226 S245E | gp226_S245E_F | gtaatcgaataccttgacgttgtgatcaaacag |
| | gp226_S245E_R | aaggtattcgattaccgagctattgtgtttcaac |
| **PCR primers used to create the plasmid for expression of the tagless version of AR9 nvRNAP core** | | |
| | *Oligonucleotide name* | *5′-3′ sequence* |
| | nvRNAP-tag-free-for-66.5 | tttaactttaagaaggagatataccatggggaaaaaattatcgttaatcgatttcaac |
| | nvRNAP-tag-free-rev-72.1 | cagcagcggtttctttaccagactcgagttatttttcatc |
| **DNA oligonucleotides in the AR9 nvRNAP promoter complex used for structure determination** | | |
| | *Oligonucleotide name* | *5′-3′ sequence* |
| | [+3;+16] non-template strand | atcacatattggag |
| | [-16;+16] template strand all U | ctccaatatgtgatataatatauuguuuattg |
| **DNA templates used to examine the dependence of *in vitro* transcription activity of the AR9 nvRNAP on the position and number of T bases in the promoter (Extended data figure 1c)** | | |
| *Name of the template in the figure* | *Oligonucleotide name* | *5′-3′ sequence* |
| all U | [+3;+16] non-template strand | atcacatattggag |
| | [-16;+16] template strand all U | ctccaatatgtgatataatatauuguuuattg |
| (-12)T | [+3;+16] non-template strand | atcacatattggag |
| | [-16;+16] template strand T(-12) | ctccaatatgtgatataatatauuguutattg |
| (-11)T | [+3;+16] non-template strand | atcacatattggag |
| | [-16;+16] template strand T(-11) | ctccaatatgtgatataatatauugutuattg |

| | | |
|---|---|---|
| (-10)T | [+3;+16] non-template strand | atcacatattggag |
| | [-16;+16] template strand T(-10) | ctccaatatgtgatataatatauugtuuattg |
| (-8)T | [+3;+16] non-template strand | atcacatattggag |
| | [-16;+16] template strand T(-8) | ctccaatatgtgatataatatautguuuattg |
| (-7)T | [+3;+16] non-template strand | atcacatattggag |
| | [-16;+16] template strand T(-7) | ctccaatatgtgatataatatatuguuuattg |
| all T | [+3;+16] non-template strand | atcacatattggag |
| | [-16;+16] template strand all T | ctccaatatgtgatataatatattgtttattg |
| **DNA templates used to examine the *in vitro* transcription activity of the AR9 nvRNAP gp226 V206G mutant (Figure 2b)** | | |
| *Name of the template in the figure* | *Oligonucleotide name* | *5′-3′ sequence* |
| all U | [+3;+16] non-template strand | atcacatattggag |
| | [-16;+16] template strand all U | ctccaatatgtgatataatatauuguuuattg |
| (-11)T | [+3;+16] non-template strand | atcacatattggag |
| | [-16;+16] template strand T(-11) | ctccaatatgtgatataatatauugutuattg |
| (-10)T | [+3;+16] non-template strand | atcacatattggag |
| | [-16;+16] template strand T(-10) | ctccaatatgtgatataatatauugtuuattg |
| all T | [+3;+16] non-template strand | atcacatattggag |
| | [-16;+16] template strand all T | ctccaatatgtgatataatatattgtttattg |
| **DNA templates used to examine the *in vitro* transcription activity of the AR9 nvRNAP gp226 Y246A and gp226 S245E mutants (Figure 2d)** | | |
| *Name of the template in the figure* | *Oligonucleotide name* | *5′-3′ sequence* |
| ds DNA | [-16;+16] non-template strand | caataaacaatatattatatcacatattggag |
| | [-16;+16] template strand all U | ctccaatatgtgatataatatauuguuuattg |
| fork DNA | [+3;+16] non-template strand | atcacatattggag |

| | | |
|---|---|---|
| | [-16;+16] template strand all U | ctccaatatgtgatataatatauuguuuattg |
| **PCR primers that were used for PCR amplification of genomic DNA fragments to examine the *in vitro* transcription activity of the $A^5$ mutant (Figure 2f)** | | |
| *Name of the template in the figure* | *Oligonucleotide name* | *5′-3′ sequence* |
| [-60;+80] DNA | P077-for-UP-60-63.4 | taatcctcctacttatctagtctataattaattgttg |
| | P077-rev-ROff-80-61.9 | attgcttcattaaacataaatgaagactc |
| [-16;+80] DNA | P077-for-UP-16-60.4 | caataaacaatatattatatcacatattggagg |
| | P077-rev-ROff-80-61.9 | attgcttcattaaacataaatgaagactc |

**Supplementary Table 2. X-ray data collection and refinement statistic.**

| | AR9 nvRNAP core native (PDB code: 7S00) | AR9 nvRNAP core thimerosal (Hg) derivative | AR9 nvRNAP core $Ta_6Br_{12}$ derivative | AR9 nvRNAP core thimerosal (Hg) derivative large unit cell | AR9 nvRNAP promoter complex native (PDB code: 7S01) |
|---|---|---|---|---|---|
| **Data collection** | | | | | |
| Beamline | APS 21-ID-G | APS 21ID-F | ALS 5.0.2 | APS 21-ID-D | APS 21-ID-D |
| Detector | Rayonix MX-300 | Rayonix MX-300 | Dectris Pilatus3 6M 25Hz | Dectris Eiger 9M | Dectris Eiger 9M |
| Wavelength (Å) | 0.97857 | 0.97872 | 1.25515 | 1.0050 | 0.91840 |
| Space group | $P2_12_12_1$ | $P2_12_12_1$ | $P2_12_12_1$ | $P2_12_12_1$ | C2 |
| Cell dimensions | | | | | |
| *a*, *b*, *c* (Å) | 112.86, 166.27, 307.22 | 113.43, 169.51, 308.30 | 112.77, 171.51, 309.76 | 171.24, 231.78, 592.45 | 176.93, 110.46, 222.38 |
| α, β, γ (°) | 90.00, 90.00, 90.00 | 90.00, 90.00, 90.00 | 90.00, 90.00, 90.00 | 90.00, 90.00, 90.00 | 90.00, 98.52, 90.00 |
| Resolution (Å) | 50.0-3.30 (3.50-3.30)* | 50.0-3.60 (3.82-3.60) | 50.0-4.53 (4.81-4.53) | 50.0-3.79 (4.02-3.79) | 50.0-3.38 (3.58-3.38) |
| $R_{merge}$ (%) | 12.3 (124.2) | 18.2 (201.2) | 20.6 (172.2) | 20.9 (138.0) | 15.1 (101.1) |
| $I / \sigma I$ | 10.07 (1.17) | 8.77 (1.13) | 6.75 (1.07) | 6.22 (1.02) | 7.91 (1.13) |
| Completeness (%) | 98. 6 (98.7) | 99.7 (98.5) | 99.6 (98.3) | 99.4 (99.7) | 95.3 (85.7) |
| Redundancy | 4.55 (4.20) | 6.95 (6.87) | 6.25 (6.44) | 6.30 (6.18) | 3.33 (3.40) |
| $CC_{1/2}$ | 99.8 (42.1) | 99.8 (54.8) | 99.7 (47.1) | 99.5 (52.4) | 99.2 (65.0) |
| **Refinement** | | | | | |
| Resolution (Å) | 49.4 – 3.30 | | | | 50.0 – 3.40 |
| No. reflections | 86,075 | | | | 56,728 |
| $R_{work}$ / $R_{free}$ | 0.2180 / 0.2580 | | | | 0.2379 / 0.2921 |
| No. atoms | | | | | |
| Protein | 33,892 | | | | 21,749 |
| Ligand/ion | 2 ($Zn^{2+}$) | | | | 1,500 (DNA) / 96 (ions) |
| Water | 0 | | | | 0 |
| *B*-factors (Å$^2$) | | | | | |
| Protein | 154.65 | | | | 126.15 |
| Ligand/ion | 154.85 | | | | 239.32 (DNA) / 157.88 (ions) |
| Water | NA | | | | NA |
| R.m.s. deviations | | | | | |
| Bond lengths (Å) | 0.003 | | | | 0.002 |
| Bond angles (°) | 0.57 | | | | 0.462 |
| Validation | | | | | |
| MolProbity score | 1.46 | | | | 1.34 |
| Clashscore | 8.48 | | | | 6.24 |
| Poor rotamers (%) | 0.03 | | | | 0.00 |
| Ramachandran plot | | | | | |
| Favored (%) | 98.30 | | | | 98.07 |
| Allowed (%) | 1.70 | | | | 1.93 |
| Disallowed (%) | 0.00 | | | | 0.00 |

*Values in parentheses are for the highest resolution shell.

### Supplementary Table 3. Cryo-EM data collection and refinement

| | AR9 nvRNAP promoter complex (EMDB-24763) | AR9 nvRNAP holoenzyme (EMDB-24765) |
|---|---|---|
| **Data collection and processing** | | |
| Magnification | 80,000 | 130,000 |
| Voltage (kV) | 300 | 300 |
| Electron exposure ($e^{-}$/Å$^2$) | 43.7 | 43.2 |
| Defocus range (μm) | −1 to −4 | −1 to −3 |
| Pixel size (Å) | 1.09 | 1.08 |
| Symmetry imposed | C1 | C1 |
| Initial particle images (no.) | 420,791 | 227,577 |
| Final particle images (no.) | 106,876 | 104,471 |
| Map resolution (Å) | 3.8 | 4.4 |
| FSC threshold | 0.143 | 0.143 |
| Map resolution range (Å) | 2.5-5.5 | 3.1-6.1 |

**Supplementary Table 4. Summary of MD simulation configurations and results.**

| Free energy term | Simulation Name | Window Number | Window Steps ($\times10^3$) | Equilibration Steps per Window ($\times10^3$) | Total Time (ns) | Energy Value (kcal/mol) |
|---|---|---|---|---|---|---|
| | DNA-RNAP Alchemical | 400 | 40 | 8 | 64 | 705.7 ± 2.3 |
| | DNA (bulk water) Alchemical | 400 | 150 | 30 | 240 | 693.0 ± 0.4 |
| *It is a sum of seven energy terms with a combined (propagated) error* | DNA-RNAP Constraint: $r$ | 20 | 600 | 120 | 48 | 0.3 ± 0.0 |
| | DNA-RNAP Constraint: $\phi$ | 20 | 60 | 12 | 4.8 | 0.3 ± 0.0 |
| | DNA-RNAP Constraint: $\theta$ | 20 | 300 | 60 | 24 | 0.9 ± 0.4 |
| | DNA-RNAP Constraint: $\chi$ | 20 | 60 | 12 | 4.8 | 0.2 ± 0.0 |
| | DNA-RNAP Constraint: $\psi$ | 20 | 60 | 12 | 4.8 | 1.0 ± 0.0 |
| | DNA-RNAP Constraint: $\xi$ | 20 | 60 | 12 | 4.8 | 0.9 ± 0.0 |
| | DNA-RNAP Constraint: RMSD | 20 | 600 | 120 | 48 | 3.3 ± 1.4 |
| | DNA (bulk water) Constraint: RMSD | 19 | 10,000 | 2,000 | 760 | 12.7 ± 0.1 |

Extended data Figure 1c

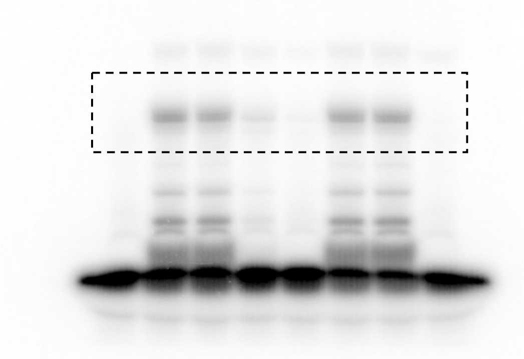

Extended data Figure 1d

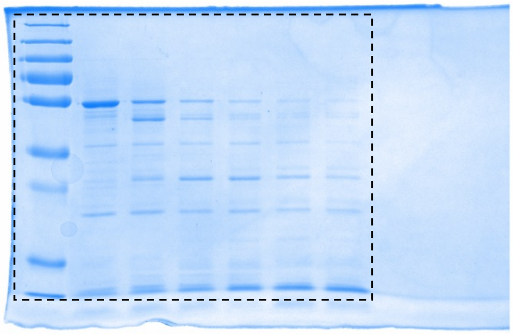

Figure 2b

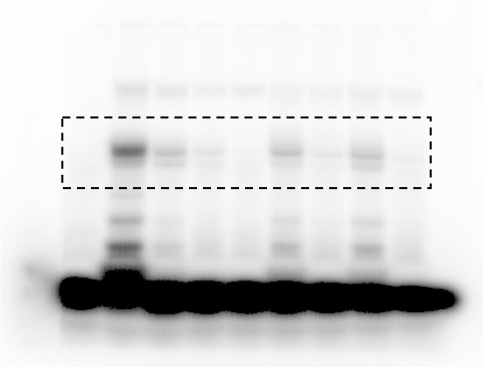

Figure 2f

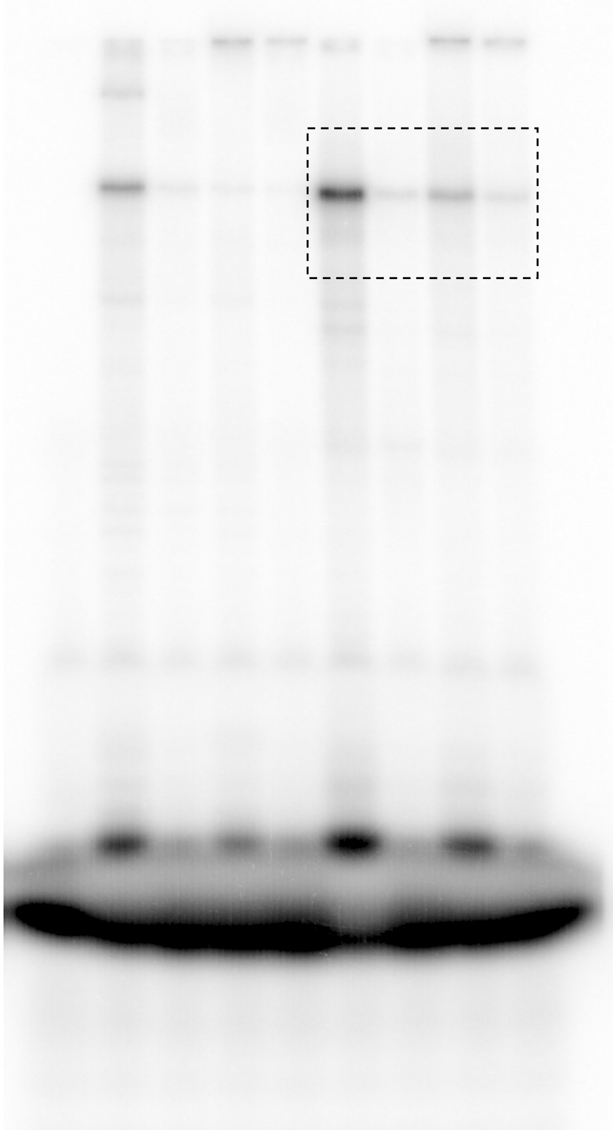

Figure 2d, top panel

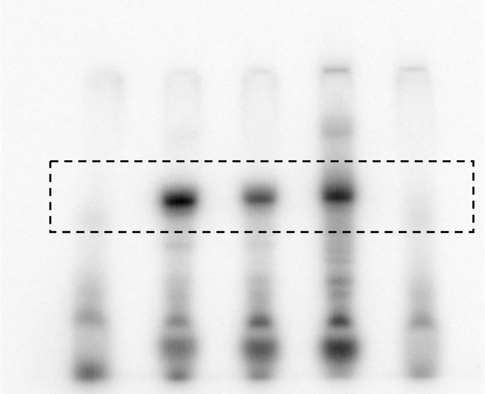

Figure 2d, bottom panel

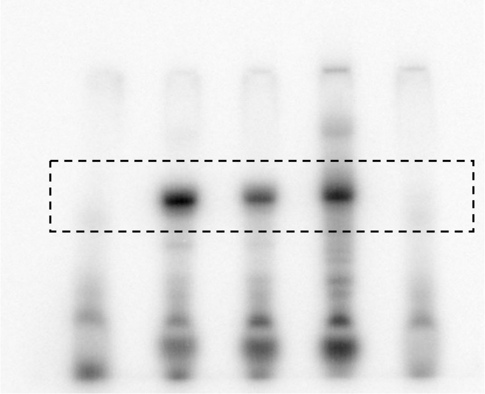

**Supplementary Figure 1. Source images of the gels used in Figures with the cropped areas marked by dashed lines.**

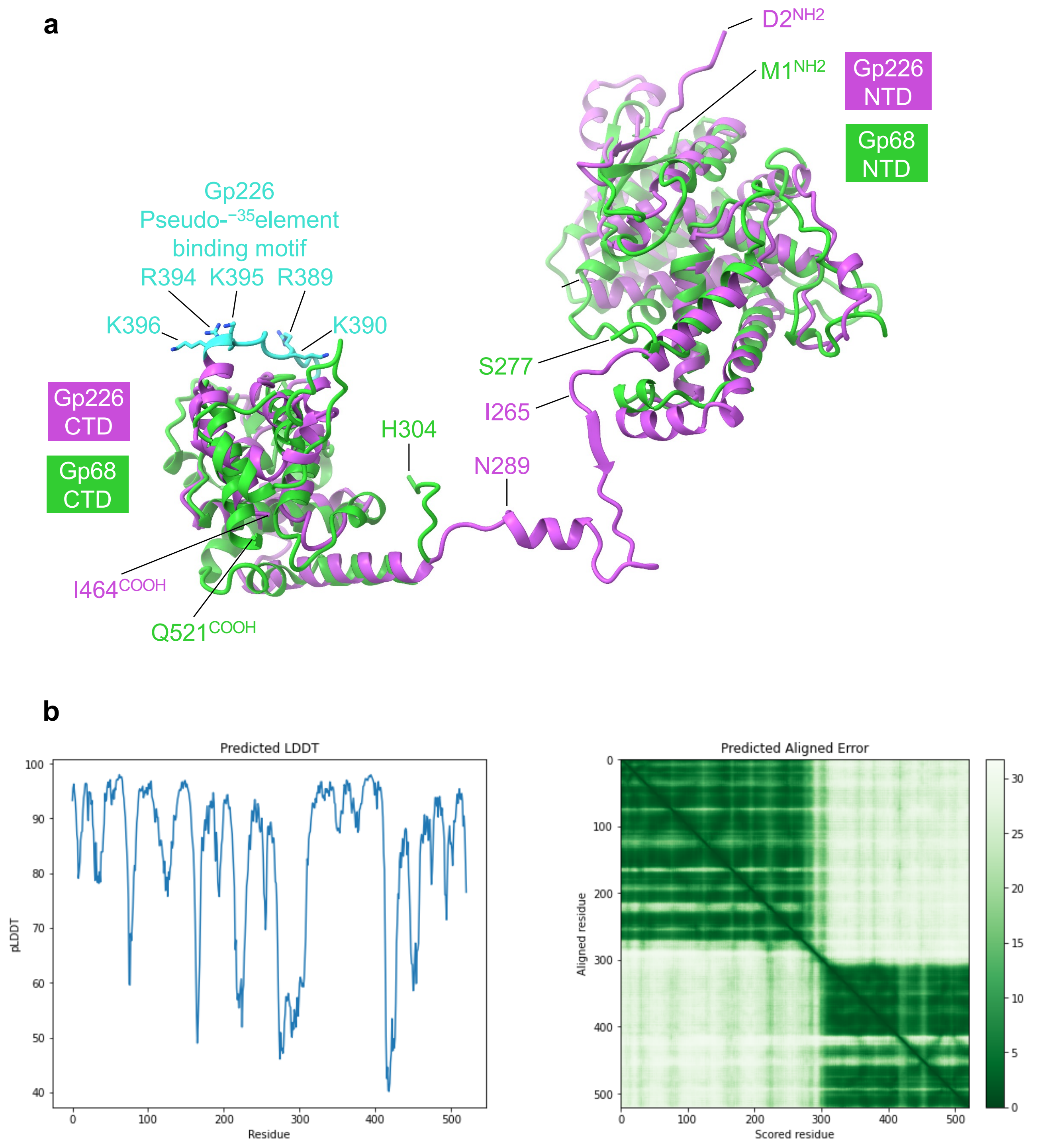

**Supplementary Figure 2. Comparison of AR9 gp226 and phiKZ gp68 structures.**
**a,** Superposition of phiKZ gp68 NTD and CTD onto gp226. The structure of gp68 NTD is predicted by AlphaFold Colab using default parameters. The structure of gp68 CTD is taken from the cryo-EM structure of phiKZ nvRNAP holoenzyme (PDB code 7OGP). The orientation of AR9 gp226 is as in **Extended Data Fig. 6**. Residue identities and numbers are given at strategic locations.
**b,** Quality factors of the gp68 model created by AlphaFold Colab. The entire sequence of gp68 was used. The accuracy of the gp68 NTD model (residues 1-277) can be evaluated by comparing the AlphaFold model of the gp68 CTD (residues 304-521) with its cryo-EM structure. The latter two can be superimposed with an RMSD of 1.2 Å for 191 equivalent Cα atoms (out of 203 ordered in the cryoEM structure).

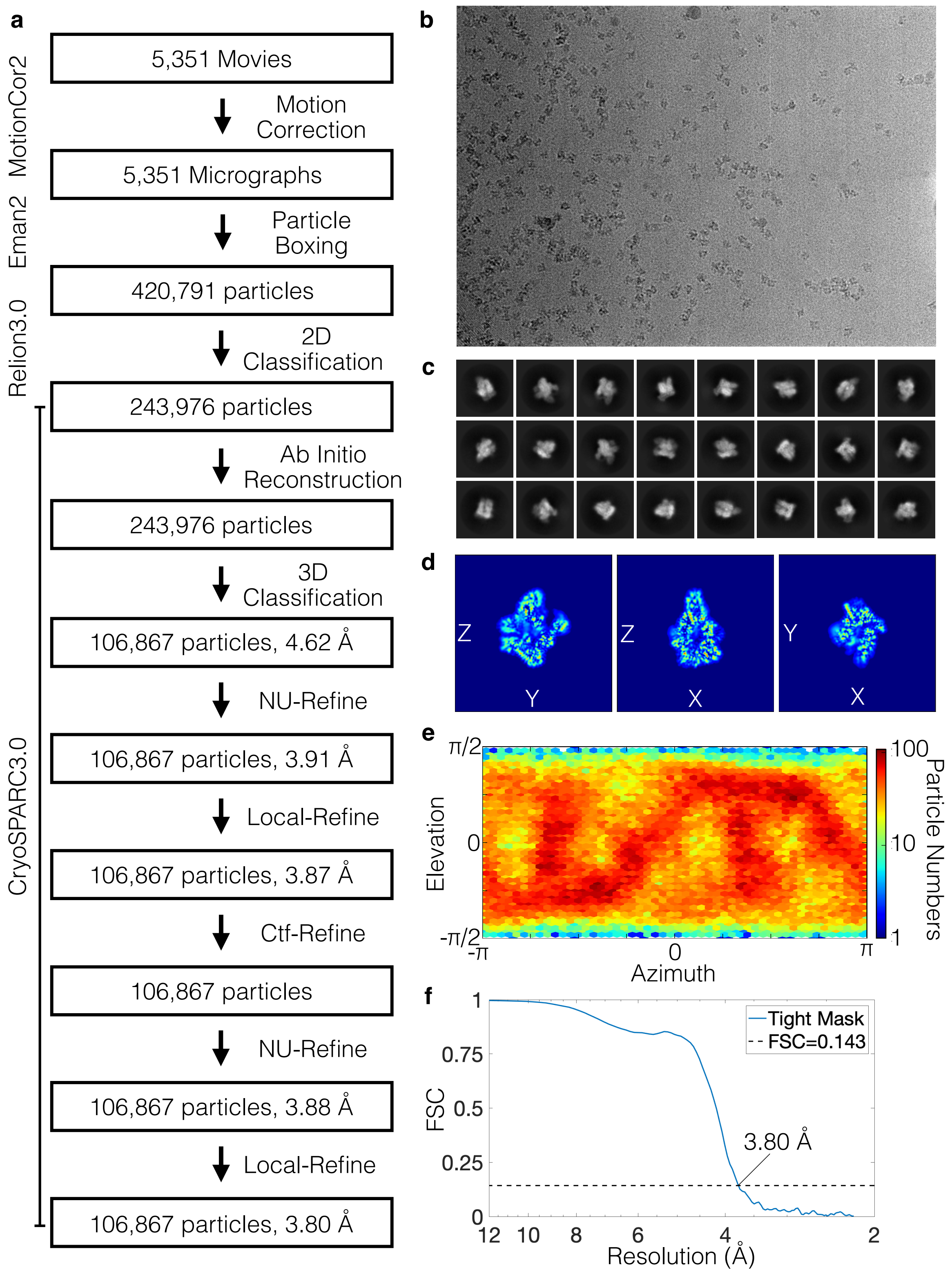

**Supplementary Figure 3. Cryo-EM image processing workflow for AR9 nvRNAP promoter complex.**
**a,** Schematic illustrating image processing jobs, software, and numbers of micrographs/particles from motion correction to the final map.
**b,** Representative micrograph after motion correction.
**c**, 2D class averages representing 244k particles.
**d**, 3 cross sections of the final map in the Z-Y, Z-X and Y-X planes.
**e,** 2D histogram (heat map) of particles based on their orientations (according to their azimuth and elevation).
**f**, FSC curve of the tightly masked map, with a nominal resolution of 3.8 Å.
